# Ubiquitin-like cGAS chain formation by a super enzyme activates anti-phage response

**DOI:** 10.1101/2022.05.25.493364

**Authors:** Yan Yan, Jun Xiao, Fengtao Huang, Bingbing Yu, Rui Cheng, Hui Wu, Xueling Lu, Xionglue Wang, Greater Kayode Oyejobi, Carol V. Robinson, Hao Wu, Di Wu, Longfei Wang, Bin Zhu

## Abstract

The cyclic oligonucleotide-based anti-phage signaling system (CBASS) is a family of defense system in prokaryotes^1, 2^. Composed of a cyclic GMP-AMP synthase (cGAS) and CBASS-associated proteins, CBASS utilizes cyclic oligonucleotides to activate antiviral immunity^3–6^. One major group of CBASS-associated proteins are homologs of eukaryotic E2 ubiquitin-conjugating enzymes. However, the function of E2 in CBASS remains elusive. Here, we report that a bacterial E2 enzyme regulates cGAS by imitating the entire ubiquitination cascade. This includes the processing of the cGAS C-terminus, conjugation of cGAS to a cysteine residue, ligation of cGAS to a lysine residue, cleavage of the isopeptide bond, and poly-cGASylation. The poly-cGASylation fully activates cGAS to produce cGAMP, which acts as an antiviral signal and leads to cell death. Our findings reveal unique regulatory roles of E2 in CBASS and provide insights into the origin of the ubiquitin system.

---

Cyclic GMP-AMP synthase (cGAS) plays a key role in mammalian cGAS-STING innate immunity^7,8^. cGAS shares a conserved catalytic domain with *Vibrio cholerae* DncV^9^ and they were classified as cGAS/DncV-like nucleotidyltransferases (CD-NTases)^3,10^. The prokaryotic CD-NTases and ancillary proteins, which are encoded by operons and involved in bacterial anti-phage defense, constitute the cyclic oligonucleotide-based anti-phage signaling system (CBASS)^2^. Certain CBASSs share evolutionary origins with the eukaryotic cGAS-STING innate immunity pathway^2,5,6^, indicating similar underlying principles between bacterial anti-phage defense and mammalian innate immunity. Over 5,000 distinct CBASSs have been identified in more than 10% of bacteria with known genome sequences^1,3,11^. About half of CBASS operons encode a CD-NTase that can generate signal molecules and an effector that induces programed cell death; while the remaining CBASS operons encode ancillary genes in addition to the CD-NTase and the effector^1,11^. Some of these ancillary components have been reported to function as a threat sensor or a regulator of bacterial defenses^4–6^; however, most of their functions and mechanisms are unclear. Among over 5,000 predicted CBASSs, 2,199 CBASSs encode homologs of ubiquitin systems^11^, which can be divided into two main categories: one category (hereafter referred to as E1E2/JAB-CBASS, Fig. 1a) contains an E1-E2 fusion protein, and a JAB deubiquitinating peptidase, but no E3 ligase^12,13^, while the other category (hereafter referred to as E2-CBASS, Fig. 1a) consisting of 616 CBASSs that only contain an E2 protein^1,11^, of which the function is elusive.

**Fig. 1.**
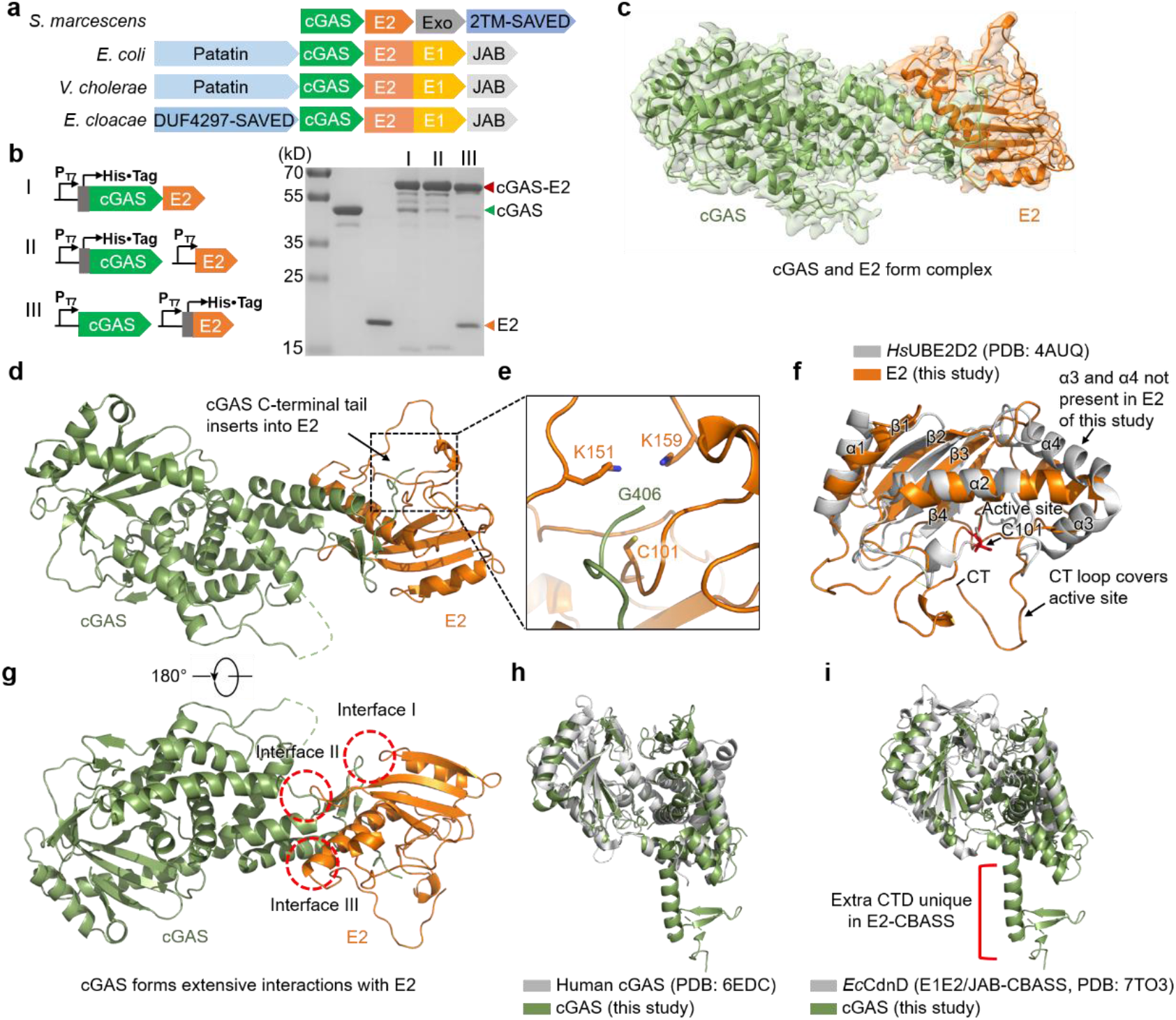
Cryo-EM structure of a cGAS-E2 complex. **a**, Domain organization of the CBASS operons from *Serratia marcescens*, *E. coli*^13^, *V. cholerae* and *Enterobacter cloacae*^12^. **b**, Left, Schematic diagram showing the constructs for co-expression of cGAS and E2 in different forms (Ⅰ–Ⅲ). Right, SDS-PAGE analysis of the purified proteins from various constructs as shown on the left. Data are representative of 3 independent experiments. **c**, Overlay of cryo-EM density and the atomic model of the cGAS-E2 complex. **d-e**, Ribbon diagram of the cGAS-E2 complex highlighting the C-terminal tail of cGAS (green) that is inserted into the E2 domain (orange). The potential interaction region (boxed) is enlarged in **e**. **f**, Superimposed structures of E2 used in this study (orange) and human E2 (gray, PDB ID 4AUQ). CT refers to the C-terminus. **g**, Ribbon diagram of the cGAS-E2 complex highlighting the interface between cGAS and E2. **h**, Superimposed structures of cGAS used in this study (green) and human cGAS (gray, PDB ID 6EDC). **i**, Superimposed structures of cGAS used in this study from E2-CBASS (green) and *Ec*CdnD^12^ from E1E2/JAB-CBASS (gray, PDB ID 7TO3). CTD refers to the C-terminal domain.

## Structure of the E2-cGAS complex

To elucidate the role of the CBASS-related E2 protein, we focused on a E2-CBASS operon from *Serratia marcescens* (Fig. 1a). The E2-CBASS operon encodes a cGAS (Extended Data Fig. 1), an E2, a putative exonuclease, and a nucleotide sensor domain (SAVED) fused to two transmembrane (TM) helices (termed 2TM-SAVED) (Fig. 1a)^1^. Reversed-phase liquid chromatography-ultraviolet (RPLC-UV) and mass spectrometry revealed that the product of cGAS is 3′,2′-cGAMP (Extended Data Fig. 1). Interestingly, when cGAS and E2 were co-expressed and co-purified, an additional band with a molecular weight of 64 kDa appears on the SDS-PAGE gel, suggesting that cGAS (46 kDa) and E2 (18 kDa) are fused (Fig. 1b). When separately-purified cGAS and E2 were mixed at 1:1 molar ratio, the linkage still formed without any cofactors in the storage buffer (50 mM Tris-HCl, pH 7.5, 100 mM NaCl, 0.1 mM EDTA, 0.1% Triton X-100, and 50% glycerol) (Extended Data Fig. 2a). This fusion of E2 and cGAS could not be separated by SDS, confirming a covalent linkage between the two.

To elucidate the structural basis of the covalent linkage between E2 and cGAS, we purified the cGAS-E2 complex and performed structural studies using cryo-EM (Extended Data Fig. 2d-f). Two cryo-EM datasets of cGAS-E2 complex were collected and processed (Extended Data Fig. 3a, b; Extended Data Table 3). The 3D reconstruction of cGAS-E2 complex was refined to 3.3 Å and 3.1 Å, respectively. Initial model of cGAS-E2 complex were generated using AlphaFold and ColabFold^14,15^ and rigid-fitted in the electron density map (Extended Data Fig. 2b, c). Although the cryo-EM density map of cGAS-E2 was anisotropic as a result of preferred orientations, it was of sufficient quality to perform model refinement guided by the initial model (Extended Data Fig. 3c-e).

The refined cryo-EM structure of cGAS-E2 is consistent with the AlphaFold prediction (Extended Data Fig. 4), with E2 binding to the C-terminal region of cGAS (Fig. 1c, d). The C-terminus of cGAS is threaded into a pocket in E2 (Fig. 1e) and in close proximity with the side chains of one Cys and two Lys residues (C101, K151, and K159). The overall structure of *S. marcescens* E2 is similar to the structure of a human E2 (UBE2D2^16^) with a root mean square deviation (RMSD) of 2.5 (Fig. 1f). C101 of *S. marcescen* E2 is located in close proximity to the active site of human E2, indicating that it is the active site cysteine. Notably, *S. marcescen* E2 has a C-terminal loop (residue 150-160) that covers the active site instead of the C-terminal α-helices (α3 and α4) found in human E2. In contrast to the E1E2/JAB-CBASS system, where cGAS interacts with dimerized E1-E2 fusion, cGAS of E2-CBASS forms extensive interactions with E2 at three interfaces in addition to the C-terminus (Fig. 1g). Although the core region of cGAS from E2-CBASS resembles human cGAS and cGAS from the E1E2/JAB-CBASS system, the C-terminal region that mediates E2 interactions is unique in cGAS from E2-CBASS (Fig. 1h, i).

## Covalent linkage between E2 and cGAS

In eukaryotes, the ubiquitin forms a thioester bond between its C-terminal Gly and a Cys in E1, E2, and E3, or forms an isopeptide bond between its C-terminal Gly and a Lys in a target protein or another ubiquitin. Similarly, the C-terminal tail of *S. marcescens* cGAS (residues 400-407) is inserted into the catalytic pocket of E2 (Fig. 1e) and the cryo-EM density of the C-terminal E407 after G406 is absent. Interestingly, G406 of cGAS is in close proximity to C101, K151, and K159 of E2, suggesting that the covalent bond between cGAS and E2 in this study is either a thioester bond between G406 of cGAS and C101 of E2, or an isopeptide bond between G406 and K151 or K159 of E2.

Alignment of the C-terminal sequences of CD-NTase homologs associated with E2 in various CBASSs showed that the glycine corresponding to cGAS G406 is highly conserved (Extended Data Fig. 5a). Deletion of the C-terminal 8 residues (ΔP400–E407) or 2 residues (ΔG406-E407) of cGAS abolished the covalent linkage between cGAS and E2, while ΔE407 had no effect. These results confirmed that G406 participates in the covalent linkage (Fig. 2a). Consistently, mutation of G406 to Val or Leu also abolished the covalent linkage, while mutation of G406 to Ala that has a shorter side chain still allowed the linkage (Fig. 2a). K401A and Y405A mutations also showed no effect (Fig. 2a).

**Fig. 2.**
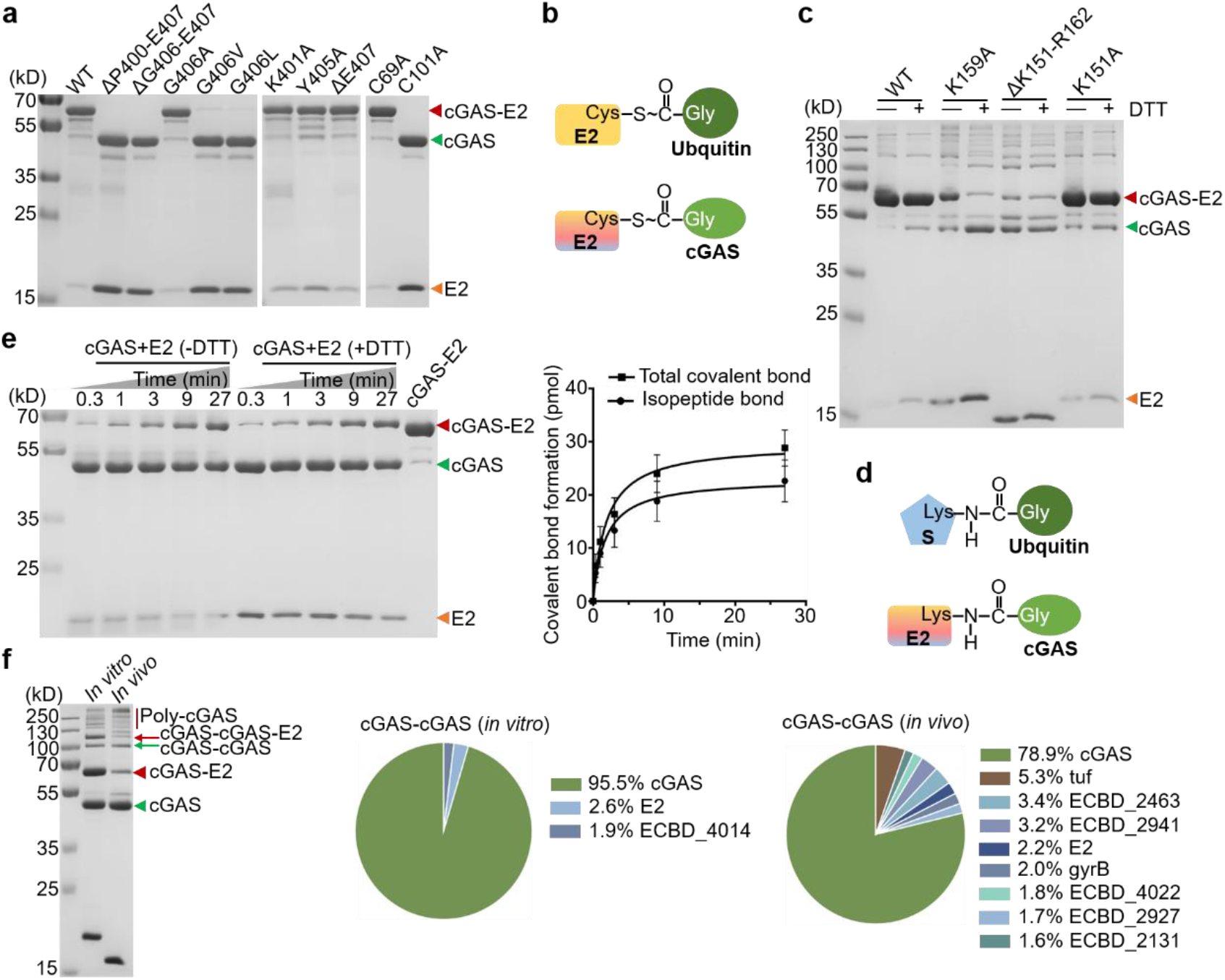
Covalent linkage between E2 and cGAS. **a**, SDS-PAGE analysis of cGAS mutants (indicated on top of gel) co-purified with E2, or E2 mutants (indicated on top of gel) co-purified with cGAS. The G406 residue of cGAS and the C101 residue of E2 were identified to be critical residues for the covalent linkage. Data are representative of 3 independent experiments. **b**, Schematic diagram showing the thioester bond between E2 and cGAS, and the thioester bond between E2 and ubiquitin. **c**, SDS-PAGE analysis of the co-purified proteins consisting of cGAS with E2 or E2 mutants in the presence (+DTT) or absence (-DTT) of 10 mM DTT (10 mM DTT was used in all following analysis). The K159 residue of E2 was identified to be the key residue for the isopeptide bonding. Data are representative of 3 independent experiments. **d**, Schematic diagram showing the isopeptide bond between E2 and cGAS, and the isopeptide bond between ubiquitin and substrate. **e**, Left, SDS-PAGE analysis showing the *in vitro* formation of the cGAS-E2 complex in the presence (+DTT) or absence (-DTT) of 10 mM DTT. Right, Rate of the total bond (thioester+isopeptide, -DTT) and isopeptide bond (+DTT) formation between cGAS and E2, as calculated based on gels in the left panel. Data are mean ± s.d. for *n* = 3 independent replicates and are representative of 3 independent experiments. **f**, Left, SDS-PAGE analysis showing the *in vivo* and *in vitro* formation of the covalent cGAS-E2 (K159A) protein. The *in vivo* formation of the covalently-linked cGAS-E2 (K159A) protein was achieved by the co-purification of His-tagged cGAS and non-tagged E2 (K159A) mutant. For the *in vitro* assembly, two proteins were mixed at a molar ratio of 1:1 overnight. Right, Mass spectrometry showing the score of proteins contained in the potential cGAS-cGAS extracted from the gel analyzing the cGAS-E2 (K159A) as shown on the left (marked by green arrow). Mass spectrometry signals corresponding to *E. coli* protein contaminants in cGAS-cGAS dimer are detailed in Extended Data Table 4.

C101 is the active site cysteine (Fig. 1f) and is conserved among prokaryotic E2 homologs (Extended Data Fig. 5b). C101A mutant failed to form the covalent complex with cGAS (Fig. 2a) in contrast to C69A that showed no effect. These results suggest that a thioester bond between cGAS G406 and E2 C101 exists (Fig. 2b).

To validate the thioester bond between G406 and E2 C101, we incubated cGAS-E2 fusion with 10 mM DTT, as the thioester bond is unstable in the presence of DTT^17^. Interestingly, the cGAS-E2 complexes are mostly resistant to DTT treatment and only a small portion was separated by 10 mM DTT (Fig. 2c), indicating that the majority of cGAS-E2 complexes are not linked by thioester bond but perhaps isopeptide bond.

To test if cGAS and E2 are linked by an isopeptide bond, we mutated K151 and K159 of E2 that are close to G406 of cGAS (Fig. 1e). Mutation of K159A but not K151A significantly reduced the cGAS-E2 complex in the presence of DTT (Fig. 2c), suggesting that the DTT-insensitive bond between cGAS and E2 is a G406-K159 isopeptide bond, which is similar to the isopeptide bond between a ubiquitin and its target protein (Fig. 2d). Interestingly, E2 K159A mutation not only reduced the DTT-insensitive isopeptide bond, but also increased the DTT-sensitive cGAS-E2 complex (Fig. 2c), indicating that the both types of covalent bonds exist and the E2 studied could catalyze the formation of both thioester and isopeptide bonds. Consistently, cryo-EM structures of cGAS-E2 exhibit heterogeneity in the E2 region, revealing conformations corresponding to both types of covalent bonds (G406-C101 and G406-K159), as depicted in Fig. 1e and Extended Data Fig. 6. Thus, bacterial E2 mimics the function of both eukaryotic E2 and E3.

As evident from the ubiquitin cascade, it is likely that the unstable thioester bond between cGAS and E2 is transient and will convert to a more stable isopeptide bond. The rates of total bond (thioester+isopeptide, -DTT, 0.63±0.17 min^-1^) and isopeptide bond (+DTT, 0.50±0.14 min^-1^) formation *in vitro* were measured and shown in Fig. 2e.

## Poly-cGASylation of cGAS by E2

Notably, the disruption of G406-K159 isopeptide bond by E2 K159A mutation not only increased the G406-C101 thioester bond, but also generated larger species insensitive to DTT (Fig. 2c, f). The molecular weight of these proteins indicates that they are polymers containing various numbers of cGAS. Either co-expressed with cGAS or mixed with cGAS *in vitro*, E2 K159A mutant generated the cGAS polymers (Fig. 2f). We excised the gel bands corresponding to the cGAS-cGAS dimer from both co-expressed and *in vitro* assembled cGAS-E2 (K159A) and analyzed them with mass spectrometry. The results showed that the majority of peptides detected were from cGAS, with minimal signals from E2 and *E. coli* protein contaminants (Fig. 2f), supporting the formation of cGAS-cGAS dimer. These cGAS polymers are insensitive to DTT, suggesting that they are linked by isopeptide bonds similar to poly-ubiquitin. Deletion of the C-terminus of E2 including K151 and K159 (ΔK151-R162) resulted in more significant poly-cGASylation, and less thioester bond (Fig. 2c).

## Cysteine protease activity of E2

The formation of either the G406-C101 thioester or the G406-K159 isopeptide bond requires the removal of the last C-terminal residue E407 of cGAS in order to expose the carboxyl group of G406. In eukaryotes, the removal of C-terminal residues of ubiquitin is carried out by specific proteases^18^. Since the single E407 is too small to observe, we constructed a cGAS-MBP by attaching a maltose binding protein to the C-terminus of cGAS to monitor the process. When cGAS-MBP was mixed with E2 *in vitro*, the presence of a protein with the same molecular weight as cGAS-E2 (64 kDa) suggested that the covalent linkage between cGAS-MBP and E2 is formed and the C-terminus of cGAS-MBP was removed (Fig. 3a). The presence of a protein with the same molecular weight as MBP further confirmed that the C-terminus of cGAS was indeed removed during the covalent bonding (Fig. 3a).

**Fig. 3.**
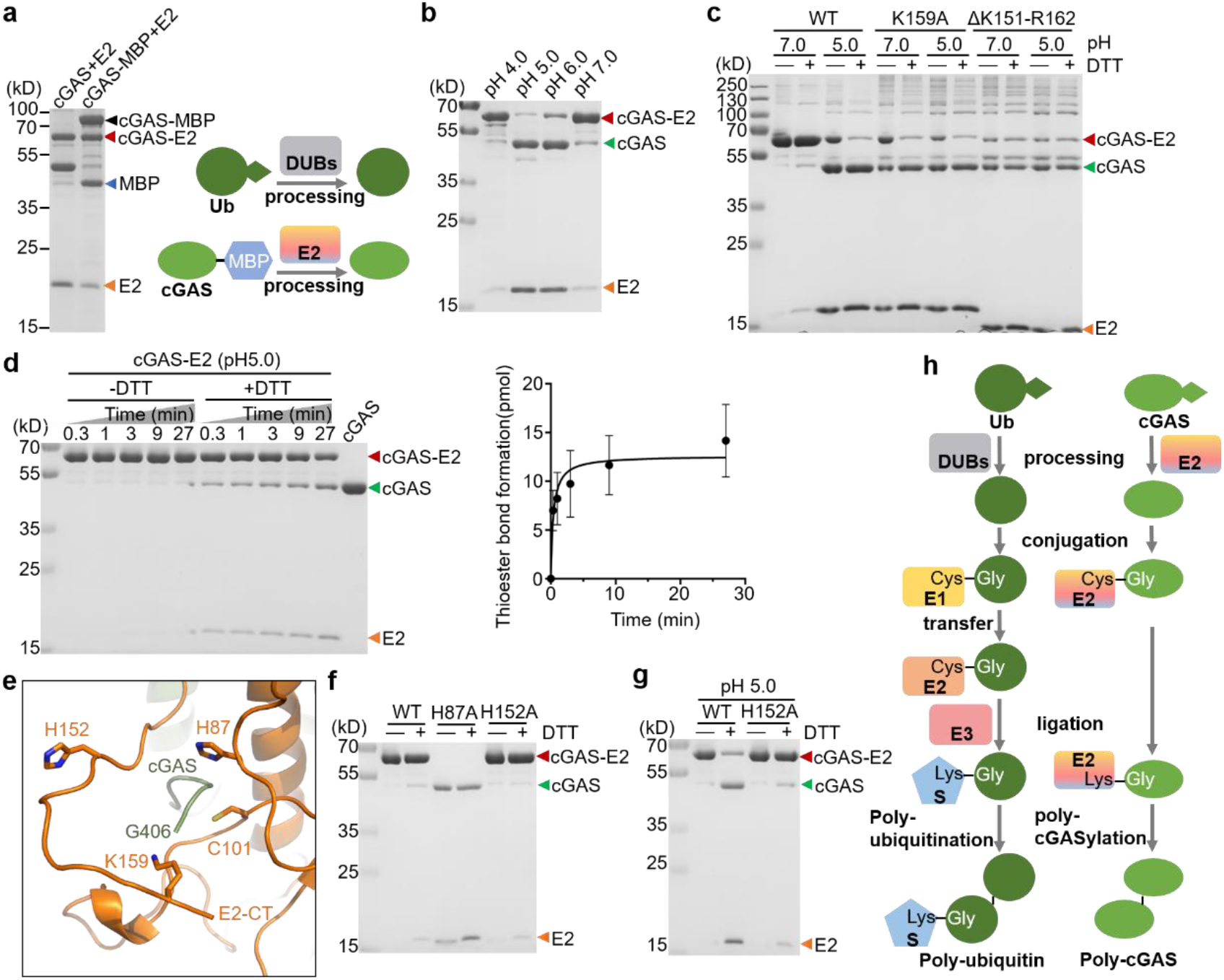
E2 is a cysteine protease. **a**, Left, SDS-PAGE analysis showing the *in vitro* processing and bonding of cGAS-MBP by E2. cGAS-MBP or cGAS and E2 were mixed at a molar ratio of 1:1 overnight. Right, Schematic diagram showed that the C-terminal cleavage of cGAS by E2 is similar to the C-terminal cleavage of ubiquitin by eukaryotic DUBs. **b**, SDS-PAGE analysis showing the low-pH-induced autocleavage of the covalent cGAS-E2 complex. The optimal pH for the autocleavage is 5.0. 10 mM DTT is presented in all reactions. Data are representative of 3 independent experiments. **c**, SDS-PAGE analysis of the low-pH-induced bond conversion in cGAS-E2 or cGAS-E2 (K159A) or cGAS-E2 (ΔK151-R162) in the presence (+DTT) or absence of DTT (-DTT). The covalent cGAS-E2 complex obtained by the co-purification of His-tagged cGAS and non-tagged E2 or E2 mutants were incubated in the reaction buffer at pH 5.0 for 30 min. **d**, Rate of the thioester bond formation in cGAS-E2 at pH 5.0 (right) calculated based on gel analysis (left). At each time interval, the amount of total bonds (thioester+isopeptide, as quantification of cGAS-E2 in the absence of DTT) subtracting the amount of isopeptide bond (as quantification of cGAS-E2 in the presence of DTT) was calculated as the amount of thioester bond. Data are mean ± s.d. for *n* = 3 independent replicates and are representative of 3 independent experiments. **e**, Magnified view of E2 showing Cys101, His87, and His152 and other key residues in the active site. E2-CT refers to the C-terminus of E2. **f**, SDS-PAGE analysis of cGAS co-purified with E2 or E2 mutants (as indicated on top of gel) in the presence (+DTT) or absence of DTT (-DTT). The H87 residue of E2 was identified to be crucial for the covalent linkage. Data are representative of 3 independent experiments. **g**, SDS-PAGE analysis of the low-pH-induced autocleavage of the covalent cGAS-E2 complex in the presence (+DTT) or absence of DTT (-DTT). The covalent cGAS-E2 complex obtained by the co-purification of His-tagged cGAS and non-tagged E2 or E2 mutants were incubated in the reaction buffer at pH 5.0 for 30 min. The H152 residue of E2 was crucial to mediate the low-pH-induced cleavage of the isopeptide bond between cGAS and E2. Data are representative of 3 independent experiments. **h**, Schematic diagram showing the eukaryotic ubiquitin pathway (left) and E2-mediated cGAS regulation (right).

The eukaryotic ubiquitin system^18^ and the E1E2/JAB-CBASS system^12,13^ both rely on deubiquitinating enzymes such as DUBs or JAB, which are mostly cysteine proteases. However, processing of cGAS requires only E2, suggesting that E2 itself may function as a cysteine protease. The C101 provides structural basis for an active site cysteine. Since low-pH-induced autocleavage is well-known for classical cysteine proteases like asparaginyl endopeptidase and cathepsin L^19,20^, we examined the effect of low-pH on the cGAS-E2 complex. Interestingly, pH 5.0 induced the cleavage of the isopeptide bond between cGAS and E2, as shown by the reduction of cGAS-E2 complex in the presence of DTT (Fig. 3b). However, in the absence of DTT, we found that like the K159A mutation, pH 5.0 also increased the thioester bond; and pH 5.0 had no effect on E2 K159A and ΔK151-R162 mutants (Fig. 3c). When the pH was adjusted from 5.0 to 8.0, the thioester bond (-DTT) between cGAS and E2 were converted back to isopeptide bond (+DTT, Extended Data Fig. 7a), confirming that the two types of covalent bond are interconvertible. The rate of thioester bond formation at pH 5.0 (0.50±0.07 min^-1^) is shown in Fig. 3d.

Another typical feature of cysteine proteases is a His residue near the catalytic Cys for proton transfer during proteolytic catalysis^21,22^. Interestingly, our structure of cGAS-E2 complex reveals two His (H87 and H152) residues that are very close to C101 (Fig. 3e). The E2 H87A mutation, like C101A (Fig. 2a), abolished the covalent linkage between cGAS and E2 (Fig. 3f). Both mutants failed to cleave the MBP of cGAS-MBP (Extended Data Fig. 7b), confirming the catalytic roles of C101 and H87 in E2 protease-like activity. The E2 H152A mutation, although showed no effect on the complex formation (Fig. 3f), significantly reduced the conversion from isopeptide bond to thioester bond at pH 5.0 (Fig. 3g), suggesting that His152 is a pH-sensor at protease active site as those reported previously^23,24^. Thus, the E2 in this study possesses both the catalytic^25,26^ and regulatory elements of proteases. The protease activity appears unique for E2 from the E2-CBASS systems, as sequence alignment showed that the histidine residues corresponding to H87 and H152 are only conserved in E2 from the E2-CBASS systems, but not in E2 from the E1E2/JAB-CBASS systems (Extended Data Fig. 7c). Also, no similar histidine residue was observed in eukaryotic E2 (Extended Data Fig. 7d). Altogether, the E2 is a “super” enzyme functionally mimicking the entire eukaryotic ubiquitination system (Fig. 3h).

## cGAS is activated by E2-mediated poly-cGASylation

Human cGAS is activated through oligomerization^27,28^. The poly-cGAS observed in this study is likely the active form of cGAS. We compared the cGAMP synthesis activities of frees cGAS, cGAS-E2, cGAS-E2 (K159A), and cGAS-E2 (ΔK151-R162). The concentration of cGAS in 4 samples were kept the same (800 nM), and the latter two samples contained more poly-cGAS (Fig. 4a). Interestingly, although the cGAS-E2 linkage inhibited the activity of cGAS, cGAS-E2 (K159A) and cGAS-E2 (ΔK151-R162) significantly increased the synthesis activity of cGAS when poly-cGAS were formed (Fig. 4a). When the concentration of ATP substrate is lowered by half, there was a more significant increase of synthesis activity by poly-cGAS (Fig. 4b), suggesting that the poly-cGASylation increased the affinity of cGAS to ATP.

**Fig. 4.**
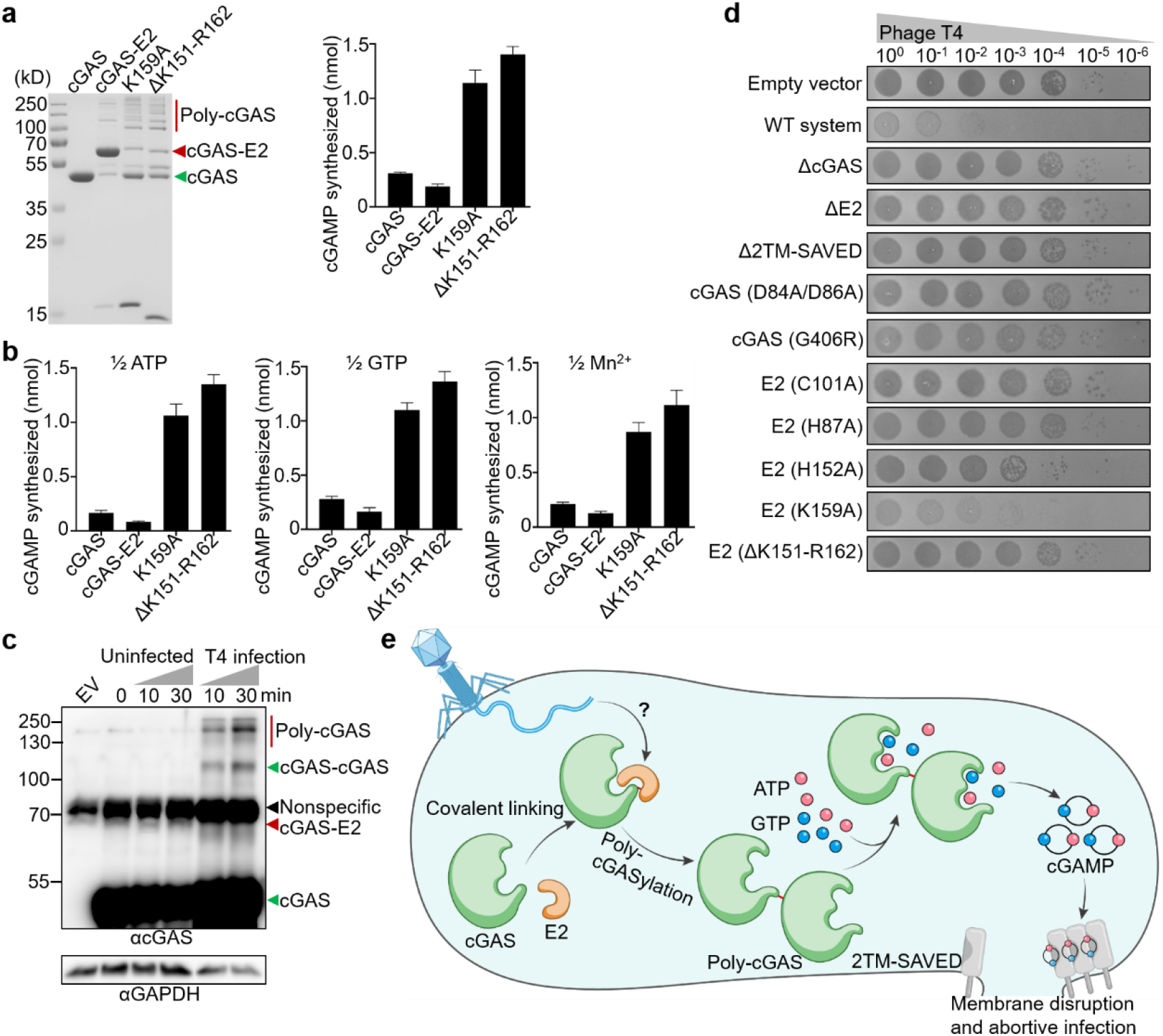
Poly-cGASylation stimulates cGAMP production and anti-phage immunity. **a**, Left, SDS-PAGE analysis of cGAS and the co-purified proteins consisting of cGAS with E2 or E2 mutants (indicated on top of gel). Right, Quantification of 3′,2′-cGAMP (based on HPLC analysis shown in Extended Data Fig. 8a) synthesized by cGAS and variants shown in left panel. Data are mean ± s.d. for *n* = 3 independent replicates and are representative of 3 independent experiments. **b**, Quantification of 3′,2′-cGAMP (based on HPLC analysis shown in Extended Data Fig. 8b) synthesized by cGAS and variants shown in (**a**), with the concentration of ATP or GTP or Mn^2+^ in the reaction reduced by half compared to those in (**a**), respectively. Data are mean ± s.d. for *n* = 3 independent replicates and are representative of 3 independent experiments. **c**, Western blot analysis with cGAS-specific antibody showing the poly-cGAS in *E. coli* BL21 expressing E2-CBASS operon during T4 phage infection. Phage T4 was applied at a multiplicity of infection (MOI) of 5. Data are mean ± s.d. for *n* = 3 biological replicates. **d**, Phage T4 infection of cells expressing empty vector, the full native E2-CBASS operon (WT system), WT system with deletion or mutations of cGAS, WT system with deletion or mutations of E2, or WT system with deletion of the effector 2TM-SAVED. Data are representative of 3 independent experiments. **e**, Model for the E2-CBASS antiviral mechanism.

Immunoblot using cGAS-specific antibody detected the poly-cGAS in the CBASS-expressing bacteria upon T4 phage infection (Fig. 4c), confirming that the phage-induced poly-cGASylation occurs *in vivo*. Recent work has demonstrated that CD-NTase+2TM-SAVED operons utilize antiviral nucleotide signals to activate CBASS TM effectors, which in turn induce inner-membrane disruption and cause cell death to limit phage replication^29^. With this effector, the antiviral signal producer cGAS, and a regulatory E2, the E2-CBASS operon confers strong immunity on *E. coli* against bacteriophage T4, as judged by plaque formation (Fig. 4d). This immunity was abolished upon: (1) deletion of either cGAS, or E2, or the effector 2TM-SAVED gene; (2) disruption of the second messenger synthesis by the cGAS D84A/D86A mutation; (3) disruption of the covalent linkage by the cGAS-G406R, E2-C101A, or E2-H87A mutation; (4) disruption of the low pH-dependent bond conversion by the E2-H152A, or E2-ΔK151-R162 mutation (Fig. 4d). Although K159 is the preferred residue for stable isopeptide bond formation, its mutation did not show significant effect on the antiviral immunity (Fig. 4d), indicating that the partial releasing of poly-cGAS by K159A does not impede the antiviral immunity.

## DISCUSSION

### Origin of the eukaryotic ubiquitin system

The ubiquitin system is a highly conserved in eukaryotes and plays a critical role in a variety of cellular processes, including protein degradation, DNA repair, and signal transduction^18,30^. The ubiquitin machinery consists of at least 5 proteins: a protease, an E1, an E2, an E3, and a target protein. We demonstrate that the bacterial E2 in our study regulates cGAS by imitating the entire ubiquitination cascade, including the cleavage of the C-terminus of cGAS or the isopeptide bond (functions like a protease), conjugation of cGAS to a cysteine residue (functions like E1 and E2), and ligation of cGAS to a lysine residue (functions like E3 and a target protein) (Fig. 3h). The multifunctional E2 provides insights into the evolution of eukaryotic ubiquitin system. Our results suggest that the ubiquitin system may have arisen from a single E2-like protein that originally regulate antiviral synthase rather than ubiquitin. Over time, the function of E2 diverged, resulting in the emergency of E1, E2, and JAB. The E1E2/JAB-CBASS^12,13^ may present this evolutionary stage. Finally, ubiquitin and E3 was incorporated for regulations of protein degradation in eukaryotes. One interesting aspect of the E2 protein described in this study is its protease activity, which appears to be unique among all known E2s to date, suggesting that this particular E2 may have a more primitive status in evolution. Our work also provides a compelling evidence for the central role of E2 in ubiquitin systems^31,32^.

### Poly-cGASylation by E2 for cGAS activation

The poly-cGASylation for cGAS activation revealed in this study is unique. However, it shares similarity with human cGAS and cGAS in the E1E2/JAB-CBASS system. The human cGAS depends on oligomerization for activation^27,28^; and bacterial cGAS in the E1E2/JAB-CBASS system is activated by conjugation to other proteins^12,13^. Activation through inter-protein interaction is likely a common feature for cGAS. The poly-cGASylation in this study was triggered by mutations of the E2 C-terminus. Although the poly-cGASylation was also observed *in vivo* upon phage infection (Fig. 4c), the exact signal from phages to E2 remains to be determined. Upon the phage signal disrupting the C-terminus of E2, the sequential poly-cGASylation and cGAMP production trigger membrane disruption and abortive infection (Fig. 4e).

## Methods

### Plasmid construction

All experiments were performed with the CBASS system from *Serratia marcescens* (NCBI RefSeq NZ_KE560335.1). For the expression and purification of cGAS (WP_016928966.1) and E2 (WP_016928967.1), their coding sequences were codon-optimized and commercially synthesized by Genscript. For the co-expression of cGAS and E2, an N-terminal 6×His-tag sequence was constructed in pET28a vector. Non-tagged expression vectors were constructed from pQE82L vector, with the T5 promoter replaced by a T7 promoter for E2. Three forms of co-expression present in Fig. 1b were assessed for cGAS and E2: (1) the gene encoding cGAS with an N-terminal 6×His-tag cloned into pET28a vector, the gene encoding E2 gene without tag cloned into pQE82L vector, the T5 promoter in pQE82L replaced with a T7 promoter; (2) the gene encoding cGAS without tag cloned into pQE82L vector and the gene encoding E2 with N-terminal 6×His-tag cloned into pET28a vector; (3) the native sequence of cGAS and E2 in a polycistronic form derived from *S. marcescens* was cloned into pET28a vector. For phage infections, the four-gene CBASS system operon from *S. marcescens* spanning a nucleotide range of 26,427–29,783 were cloned into plasmid pQE82L. Details on the cloning of cGAS and E2 mutants are listed in Extended Data Table 1 and 2, respectively. Plasmids generated in this study were constructed by Gibson from synthesized genes with ≥18 base pairs of homology flanking the insert sequence using ClonExpress II One Step Cloning Kit (Vazyme Biotech). Plasmids were transformed into *E. coli* DH5α cells and confirmed by Sanger sequencing.

### Protein expression, co-expression and purification

Plasmids for protein expression were transformed into *E. coli* BL21 (DE3). To determine the interaction between cGAS and E2, they were co-transformed into *E. coli* BL21 (DE3) and selected with their respective antibiotic marker. Bacterial cells were cultivated as a 4 mL starter culture in LB broth overnight at 37 °C with shaking at 220 rpm. The cultures were then transferred into 200 mL LB broth and cultivated for approximately 3 h until the optical density at 600 nm (OD_600_) reached 0.8–1.0. After cooling down to room temperature, 0.25 mM IPTG was added to the cultures, and cultivation was continued overnight at 16 °C.

Cultures were collected and lysed by sonication in Lysis buffer (20 mM Tris-HCl, pH 8.0, 300 mM NaCl, 20 mM imidazole), and the lysates were clarified by centrifugation at 21,000 × *g*, 4 °C for 1 h and filtered through a 0.45-μm filter. Proteins were purified by affinity chromatography using Ni-NTA resin (Qiagen) and a gravity column (Biorad). Ni-NTA resin was pre-equilibrated with Lysis buffer, bound to target proteins, washed with Wash buffer (20 mM Tris-HCl, pH 8.0, 300 mM NaCl, and 50 mM imidazole for cGAS and its mutants; 20 mM Tris-HCl, pH 8.0, 300 mM NaCl, and 50, 80, or 100 mM imidazole for E2, its mutants and the co-expressed proteins) and eluted with Elution buffer (20 mM Tris-HCl, pH 8.0, 300 mM NaCl, and 100 mM imidazole for cGAS, its mutants and the co-expressed proteins; 20 mM Tris-HCl, pH 8.0, 300 mM NaCl, and 200 mM imidazole for E2 and its mutants). Proteins were filter-concentrated using centrifugation and a 10-kDa or 30-kDa cut-off column (Millipore Sigma) and dialyzed at 4 °C for ∼24 h against Dialysis Buffer (50 mM Tris-HCl pH 7.5, 100 mM NaCl, 0.1 mM EDTA, 0.1% Triton X-100 and 50% glycerol). Proteins were analyzed by SDS-PAGE with Coomassie blue (Bio-Rad) staining.

### Structure prediction

ColabFold was used to predict the structure of the cGAS-E2 complex. Multiple sequence alignment (MSA) was performed using MMseqs2 (UniRef + Environmental). Structural prediction was performed using AlphaFold on a virtual cluster node with a Nvidia Tesla P100 GPU.

### Cryo-EM sample preparation and data collection

cGAS-E2 complex was further purified by size-exclusion chromatography on a HiLoad 16/600 Superdex 200 pg column (Cytiva) in TBS buffer (20 mM Tris-HCl, pH 8.0, 100 mM NaCl). Proteins were concentrated to 3-7 mg/mL, flash-frozen in liquid nitrogen, and stored at −80°C.

A 3 µL-drop of cGAS-E2 sample at 0.7 mg/mL was applied to glow-discharged copper Quantifoil R1.2/R1.3 grids, and blotted for 3 s in 100% humidity at 8 °C, then plunged into liquid ethane using a Vitrobot Mark IV. All grids were screened using a Glacios microscope. The data were collected on a 300 keV Titan Krios G4 microscope equipped with a Gatan K3 direct electron detector and a Gatan Quantum energy filter at the Cryo-EM Unit of Core Facility of Wuhan University.

For the initial dataset, 5,883 movies were collected in super-resolution mode, with 40 frames per movie, 1.61 s exposure time, 50 e-/Å2 accumulated dose, and 0.335 Å pixel size. For the final dataset, cGAS-E2 sample concentration was increased to 3.2 mg/mL and UltrAuFoil grids was used instead for plunging. 4,752 movies were collected in super-resolution mode, with 40 frames per movie, 2.16 s exposure time, 70 e-/Å2 accumulated dose, and 0.335 Å pixel size.

### Cryo-EM data processing and model building

For dataset I, movies were binned 2x (pixel size 0.67 Å) and motion-corrected using patch motion correction in CryoSPARC^33,34^, followed by patch contrast transfer function (CTF) estimation performed in the same software. Particles were picked using blob picker, binned 2x (pixel size 1.34 Å) and subjected to two rounds of 2D classifications in CryoSPARC with number of online-EM iterations set to 40 and batch size per class set to 400. 869,610 particles representing the binary complex were selected, and were used for ab-initio 3D reconstruction and heterogeneous refinements (Extended Data Fig. 3a). One class out of four were selected for 3D refinement using homogeneous refinement in C1 symmetry, resulting in a 3.8 Å cryo-EM density map of cGAS-E2. After global CTF refinement, a soft mask excluding the flexible terminal loops of cGAS-E2 complex was generated and used for local refinement in CryoSPARC. Particles were unbinned and used for another round of local refinement, and the final map resolution of cGAS-E2 complex was 3.35 Å.

The data processing procedures of dataset II were similar to dataset I, with a few differences (Extended Data Fig. 3b). (1) A general Topaz^35–38^ model was trained with the particles from the initial dataset and used for particle picking. Blob picker was also used for picking and after the first round of 2D classification, particles picked from the two pickers were merged and duplicate particles were removed. (2) Heterogeneous refinement (3D classification) was performed on 512,003 particles from 2D classification using the low-passed 3.35 Å map as initial models. (3) The final cryo-EM map of cGAS-E2 complex reached a resolution of 3.0 Å. Some degree of preferred orientation was present in both of the datasets, resulting in anisotropic maps (Extended Data Fig. 3c, d). DeepEMhancer^39^ was used for post-processing to improve the anisotropic map, which helped with the interpretation of the map. The initial model of cGAS-E2 was generated by rigid body fitting of the predicted cGAS-E2 structure into the cryo-EM maps using Chimera^40^. Regions with no electron densities were removed from the model, including residues 1-4, 191-210, 389-396 from cGAS and 1-3 from E2. Linkage between cGAS G406 and E2 K159 or C101 were modeled based on the densities with local adjustments. Initial B-factors for all the atoms of the model were set to 94.7 based on the Guinier Plot from final local refinement job. Inspection, model building, and manual adjustments were performed in Coot^41^ and real-space refinements were carried out in Phenix^42,43^. Representations of cryo-EM densities and structural models were generated using ChimeraX^40^ and PyMOL^44^.

### *In vitro* bonding assay

To reconstitute the covalent attachment of cGAS to E2 *in vitro*, cGAS and E2 or their mutants were mixed in 50 mM Tris-HCl (pH 7.0), unless otherwise noted. The mixture was incubated at 4 °C for the indicated periods. The reaction was quenched by adding SDS sample buffer with DTT (Solarbio Life Sciences) or without DTT (Biosharp Life Sciences) and heating at 95°C for 5 minutes. Proteins in the solutions were separated by SDS-PAGE in the absence of reducing agent and stained with Coomassie blue. GraphPad Prism was used for data analysis.

### *In vitro* cleavage assay

To examine the effect of low-pH on the cGAS-E2 complex, cGAS-E2 complex or its mutant was incubated in citrate buffer with different pH (Shyuanye) at 4 °C for the indicated periods. The reaction was quenched by adding SDS sample buffer with DTT (Solarbio Life Sciences) or without DTT (Biosharp Life Sciences) and heating at 95°C for 5 minutes. Proteins in the solutions were separated by SDS-PAGE in the absence of reducing agent and stained with Coomassie blue. GraphPad Prism was used for data analysis.

### HPLC analysis of enzymatic reactions

CD-NTase reactions were performed essentially as previously described^3^. Briefly, 20-μL reactions contained 2 μM enzyme and 250 μM NTPs (New England BioLabs) in reaction buffer with 50 mM Tris-HCl (pH 7.5), 100 mM NaCl, 2.5 mM MnCl_2_/MgCl_2_, and 1 mM DTT, unless otherwise noted. Reactions were incubated at 37 °C for the indicated periods, inactivated at 80 °C for 10 min, and centrifuged for 15 min at 21,000 × *g* to remove precipitated protein. Reaction products were analyzed by HPLC with a ZORBAX Eclipse XDB-C18 (4.6 × 250 mm) column and an Agilent 1260 Infinity II Series LC system. Next, 10 μL of the reaction product was injected into the column and eluted with solvent A (methanol, sigma) and solvent B (20 mM ammonium acetate, sigma) at a flow rate of 1 mL/min using the following linear gradient: 0–5 min, 5% A; 5–15 min, 5–100% A; 15–20 min, 100% A. The column was re-equilibrated for 5 min at 5% A. For analyzing the reaction products treated with CIP or nuclease P1, 20 μL of the reaction product was treated with 1 μL of CIP or nuclease P1 (New England BioLabs) at 37 °C for 1 h and inactivated at 80 °C for 10 min prior to centrifugation and analysis by HPLC.

### Western blots

Rabbit cGAS polyclonal antibody was generated by a commercial vendor (AtaGenix) using a purified cGAS antigen. Serum was used at 1:1,000 for cGAS immunoblot detection. GAPDH antibody (ab125247, Abcam) was used at 1:1,000 for use as a loading control.

For whole-cell lysate analysis, *E. coli* BL21 carrying the indicated plasmid were grown to mid-logarithmic phase and protein expression was induced by 0.2 mM IPTG. Cells were infected with phage T4 for the indicated periods at an MOI of 5. Cells were lysed by FastBreak™ cell lysis reagent (Promega) at room temperature for 15 min, followed by centrifugation at 4 °C to remove cellular debris. 80 μL soluble lysates were then mixed with 20 μL 5× SDS sample buffer with DTT, heated and concentrated at 100 °C for 30 min followed by a 5-min centrifugation at 21,000× g. Proteins in the solutions were separated by SDS-PAGE, then transferred to PVDF membranes (Millipore Sigma) charged in methanol. Membranes were blocked in nonfat dry milk for 1 h at 24 °C, followed by incubation with primary antibodies diluted in antibody dilution buffer at 1:1,000 overnight at 4 °C. PVDF membranes were then incubated with the HRP-conjugated goat anti-rabbit IgG (SA00001-2, Proteintech) or HRP-conjugated goat anti-mouse IgG (SA00001-1, Proteintech) at 1:5,000 dilution in TBST for 1 h at 24 °C and detected with an ECL Chemiluminescent Substrate Kit (YEASEN) and Amersham Imager 680 (GE Health).

### Plaque assays

Phages were propagated by infecting exponentially-growing *E. coli* B culture in LB medium at an MOI of 1:100–1:1,000, incubated at 37 °C with shaking (200 rpm) until clearing of the culture, the lysate was centrifuged and the supernatant was filter-sterilized through a 0.45-μm filter.

Bacteriophage infection assay was determined using the small drop plaque assay method^45^. The E2-CBASS operon and its mutants, as indicated, were cloned into the pQE82L vector and transformed into *E. coli* BL21, respectively. Cells containing an empty vector (pQE82L) were used as control. A single bacterial colony was picked, respectively, and grown in LB broth at 37 °C to an OD_600_ of ∼0.2. E2-CBASS operon expression was induced through 50 μM IPTG, followed by further growth for 1 h to an OD_600_ of 0.6–0.7. 800 μL of cells were mixed with 25 mL of LB with 0.75% agar containing appropriate antibiotic selection and IPTG (50 μM) and the entire sample was poured onto plates. The phage stock was 10-fold serially diluted and 4-μL drops of diluted phage lysate were placed on the solidified agar. The plates were then incubated at 37 °C for 16–18 h before imaging.

### Data analysis

Data analysis was performed using Microsoft Excel 2019, OriginPro version 9.1 and GraphPad Prism version 8.0.2. Graph plotting and statistical analysis was performed using GraphPad Prism version 8.0.2.

## Acknowledgements

We thank Profs. En-Duo Wang, Wei Yan, Hongyu Hu for their advice; Prof. Ping Xu for providing materials; Xuemei Li for technical supports; and all lab members for helpful discussions. We thank Danyang Li and Yi Zeng of the Core Facility of Wuhan University for their assistance with cryo-EM grid screening and data collection. This work was supported by National Natural Science Foundation of China (grant 32150009 and 31870165 to B.Z., 31900032 to F.T.H.) and Fund from Science, Technology and Innovation Commission of Shenzhen Municipality (grant JCYJ20210324115811032 to B.Z.). Funding for open access charge: National Natural Science Foundation of China.

## Author contributions

F.T.H. and B.Z. conceived the project. Y.Y., J.X., F.T.H., D.W., L.F.W. and B.Z. designed the experiments. Y.Y., J.X. and D.W. carried out the experiments. J.X. and L.F.W. determined and analyzed structures. Y.Y., F.T.H., G.K.O., C.V.R., H.W., D.W., L.F.W. and B.Z. analyzed the data and wrote the manuscript. All authors discussed the results and contributed to the final manuscript.

## Competing interests

The authors declared that they have no conflicts of interest to this work.

## Additional Information

**Supplementary Information** is available for this paper.

**Correspondence and requests for materials** should be addressed to Bin Zhu.

**Peer review information** *Nature* thanks the anonymous reviewers for their contribution to the peer review of this work.

**Reprints and permissions information** is available at http://www.nature.com/reprints.

## Extended data

**Extended Data Table 1.**
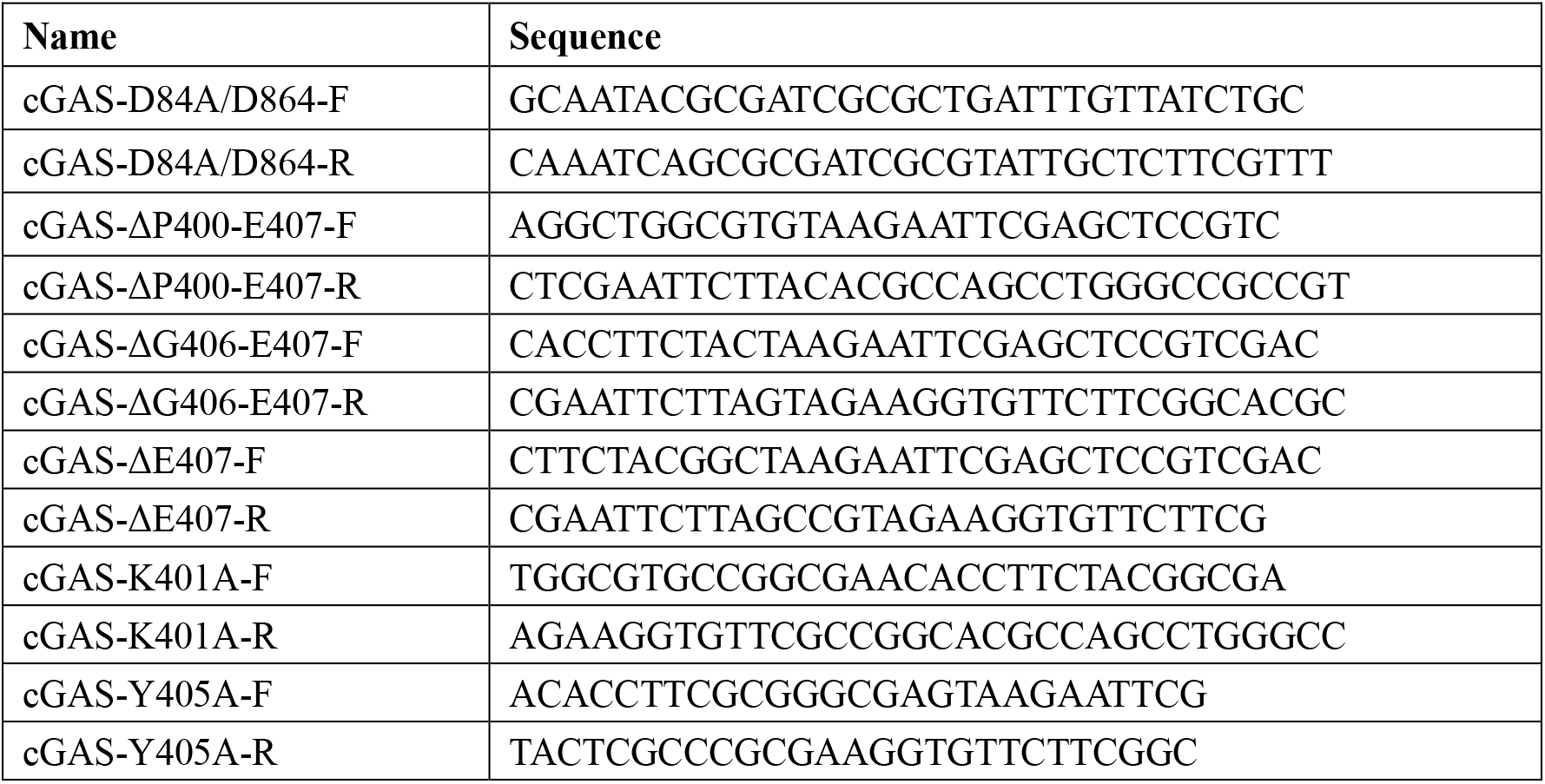
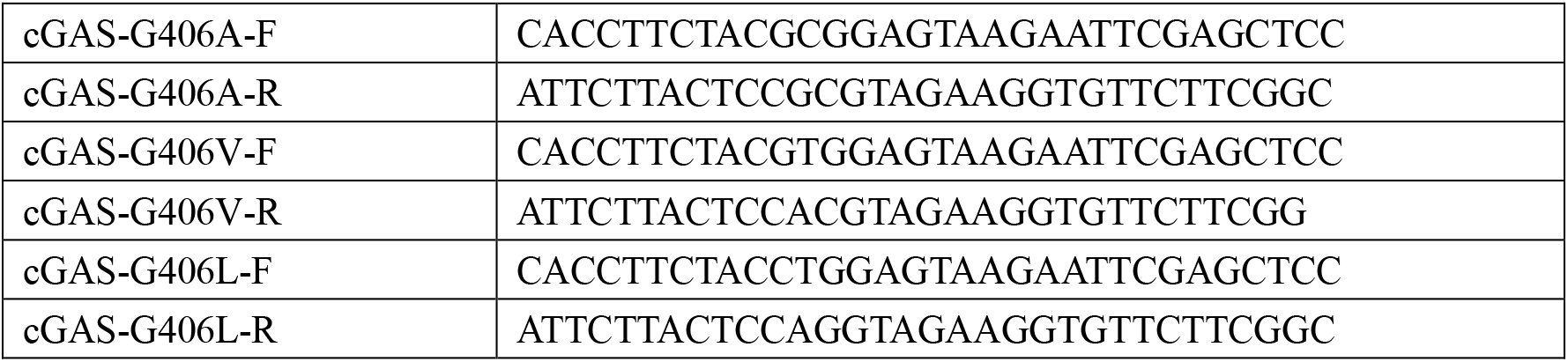
Primers for construction of cGAS mutations.

**Extended Data Table 2.**
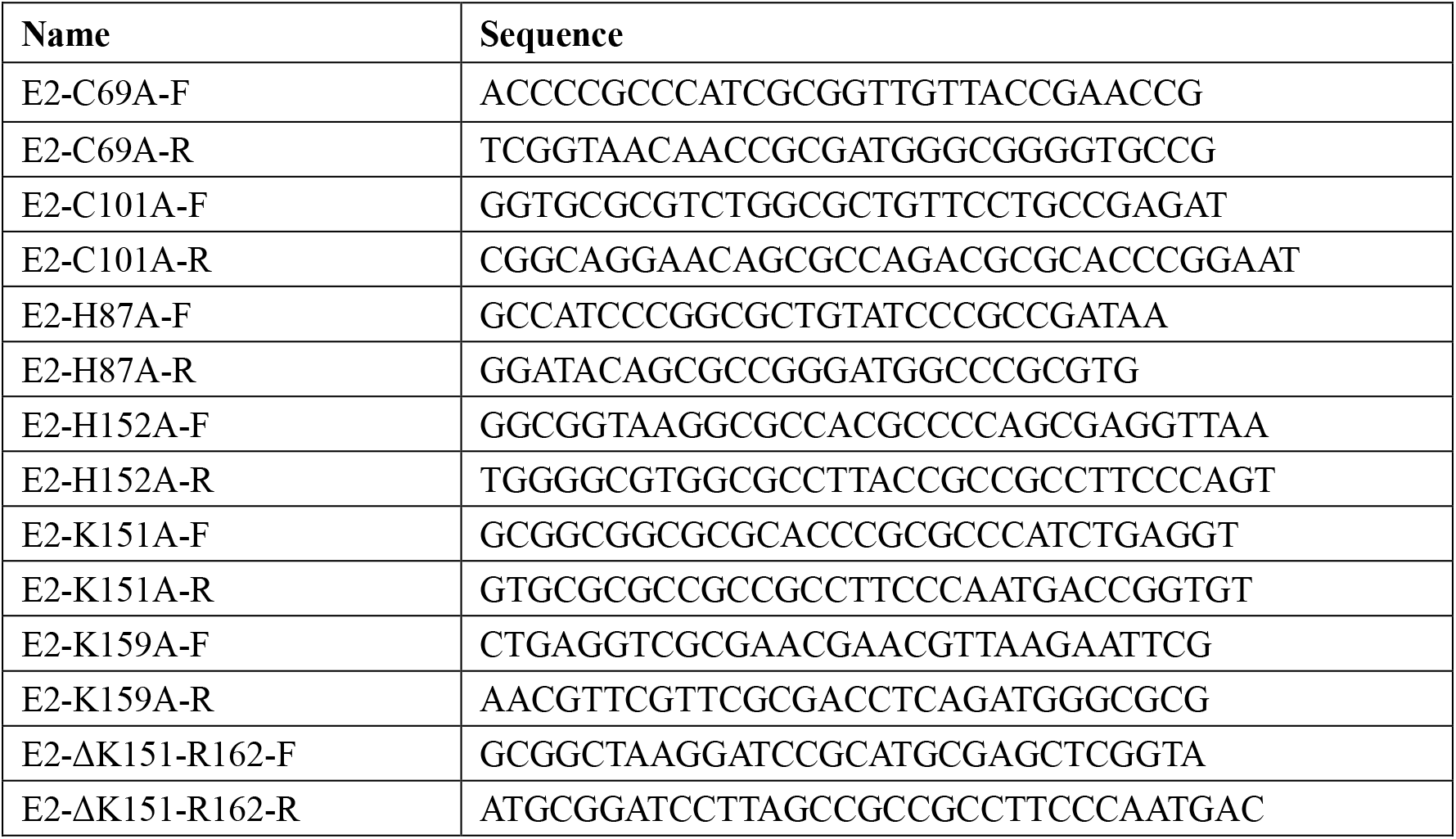
Primers for construction of E2 mutations.

**Extended Data Table 3.**
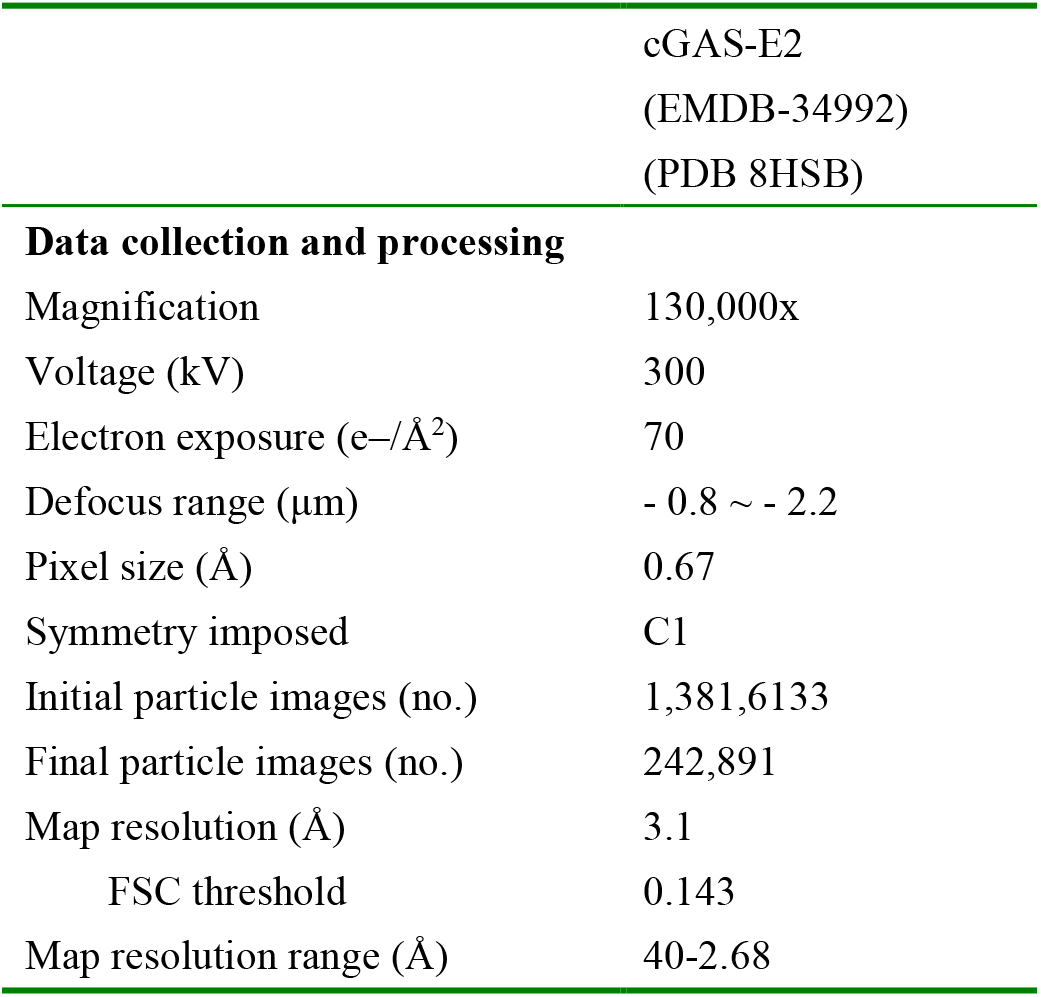
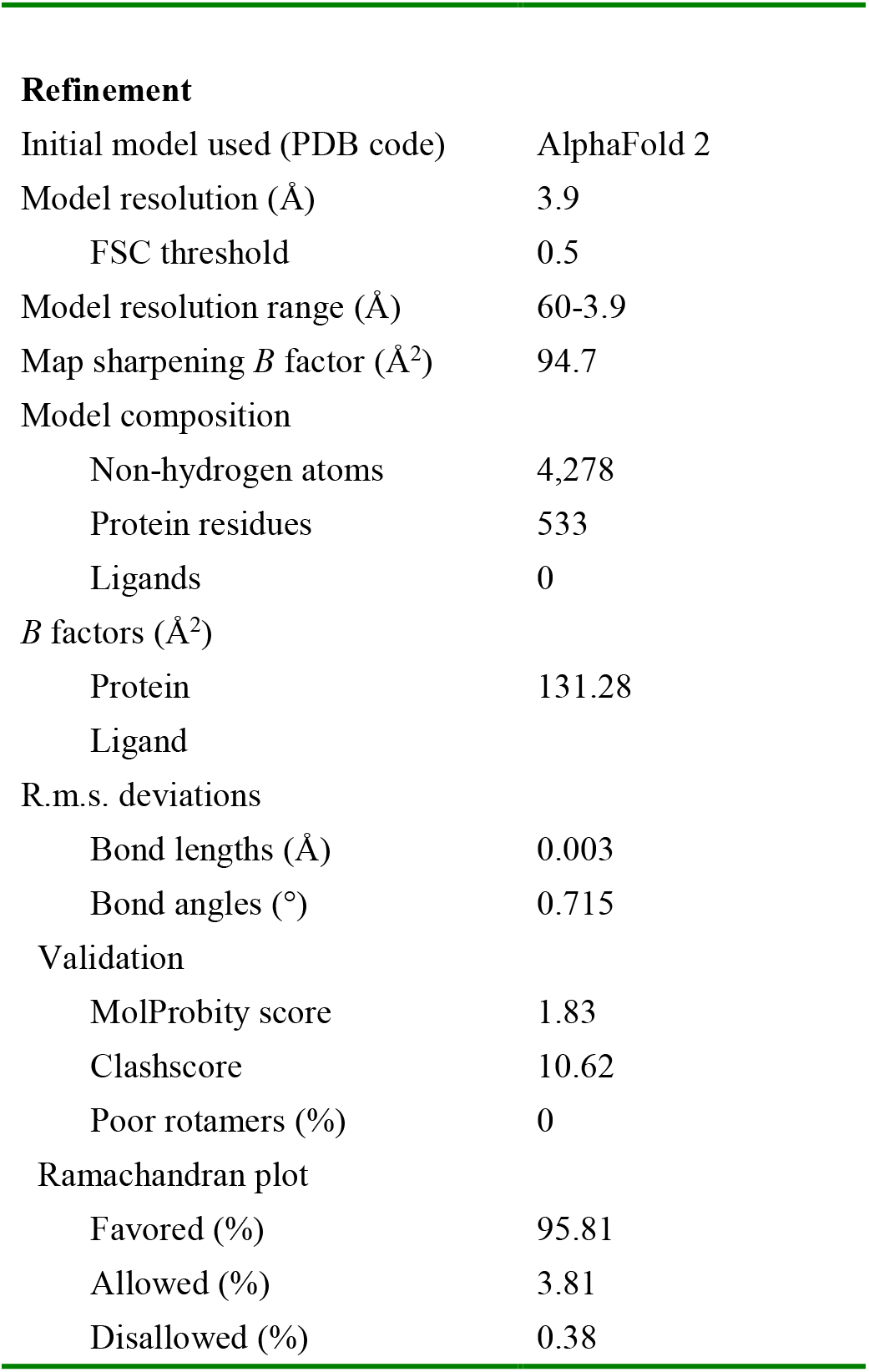
Cryo-EM data collection, refinement and validation statistics.

**Extended Data Table 4.**
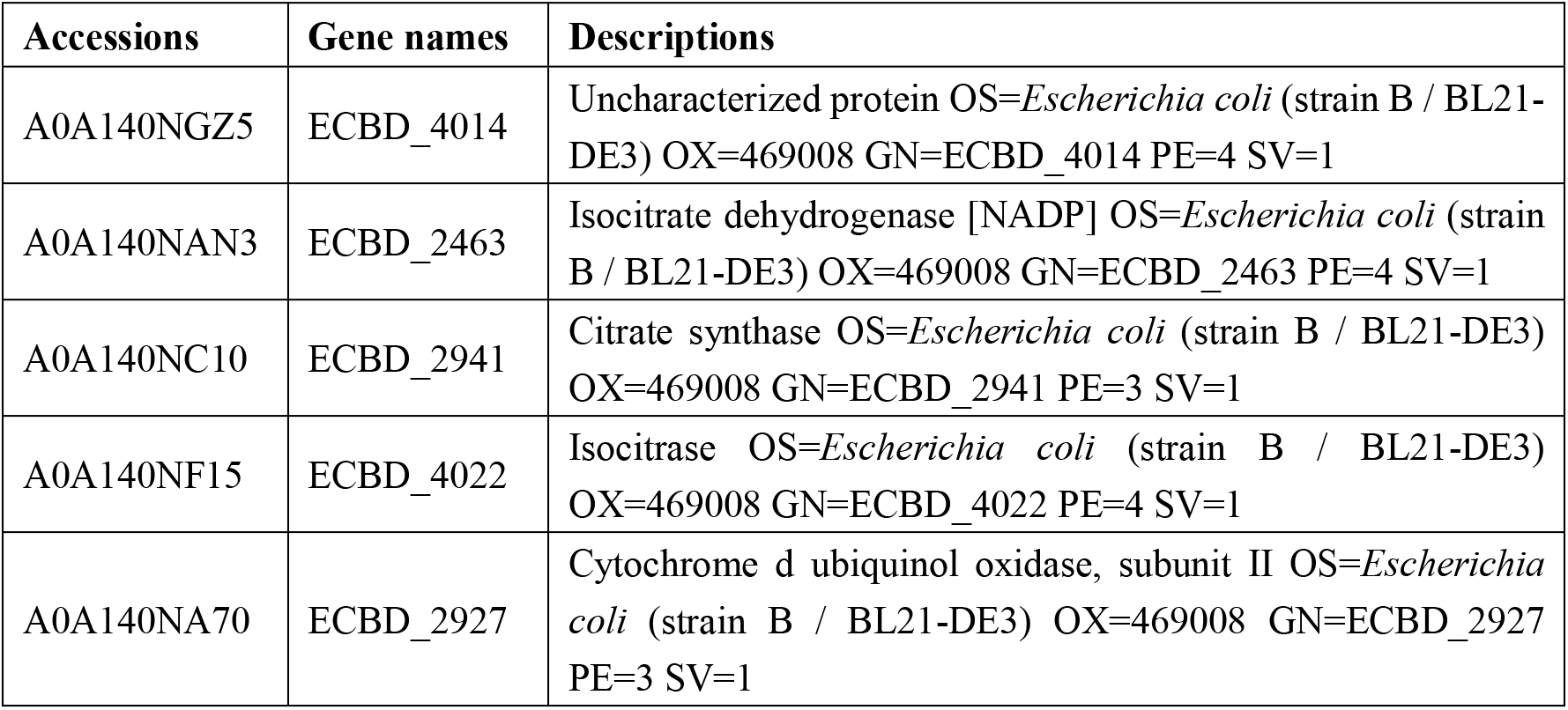
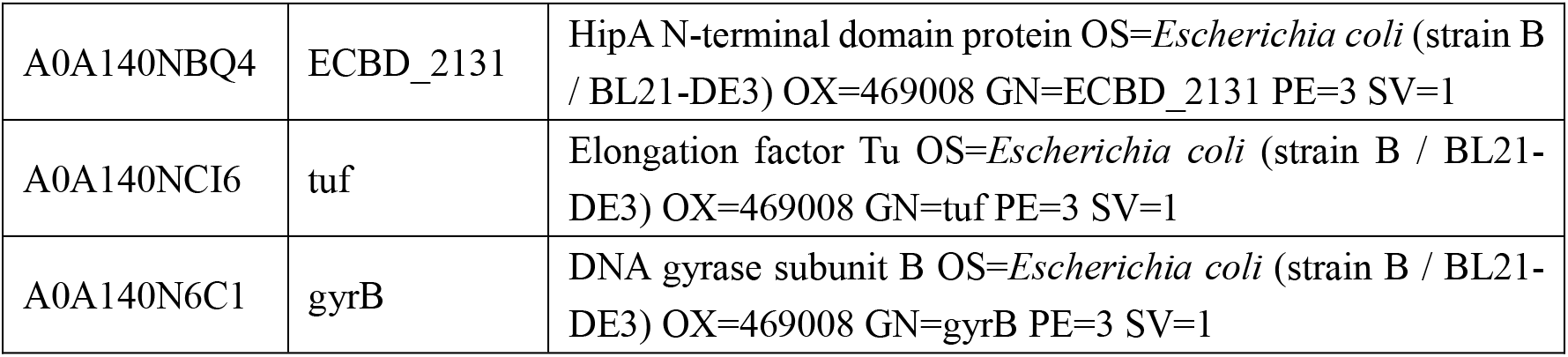
*E. coli* proteins identified in cGAS-cGAS sample by mass spectrometry.

**Extended Data Fig. 1.**
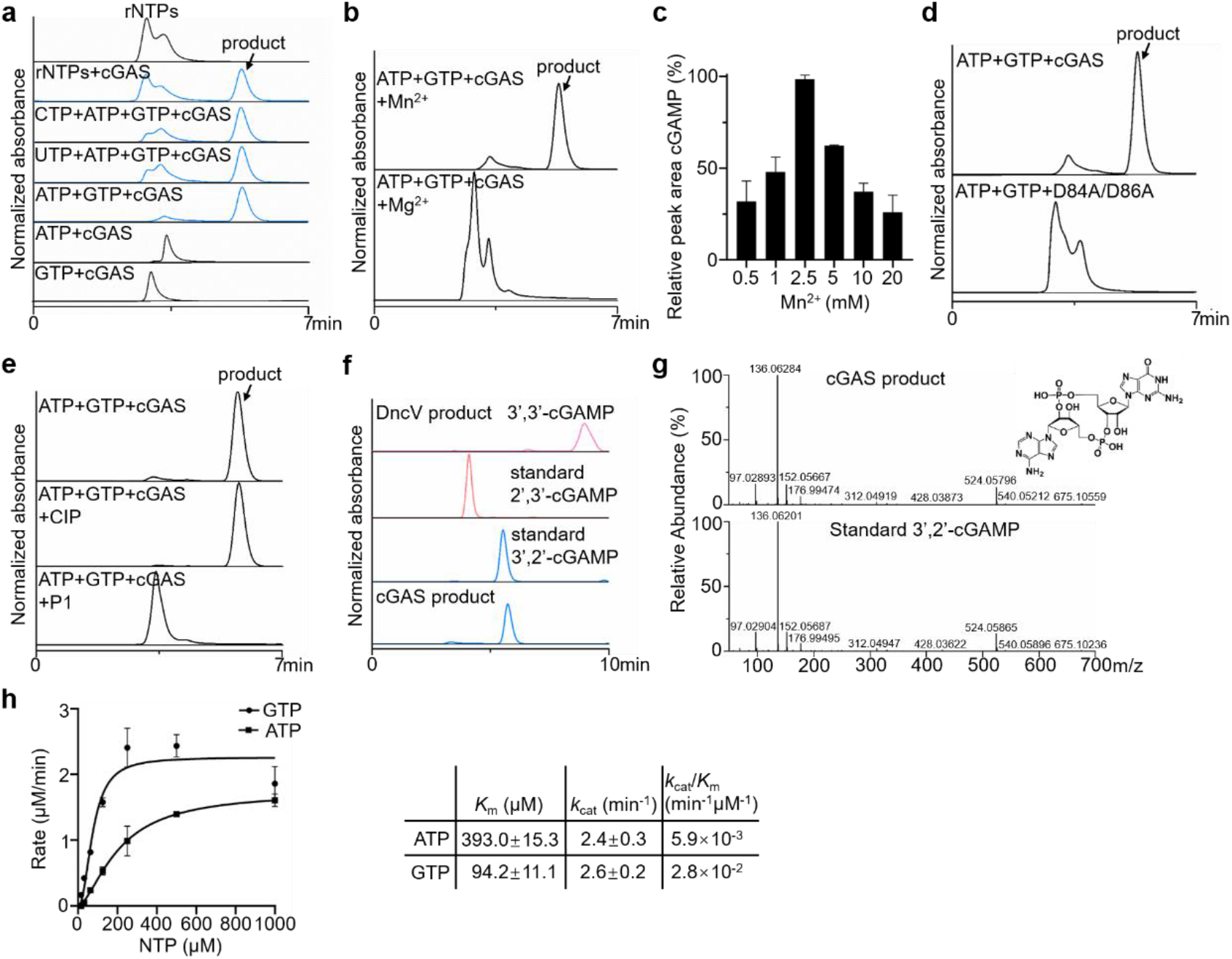
*In vitro* characterization of the enzymatic activity of cGAS. **a**, HPLC analysis of nucleotide second messenger synthesis by cGAS using various nucleotide substrates. cGAS synthesizes a product only in the presence of both ATP and GTP. Data are representative of 3 independent experiments. **b**, cGAS synthesis activity is dependent on Mn^2+^. Data are representative of 3 independent experiments. **c**, Optimal Mn^2+^ concentration for cGAS. The products synthesized at various Mn^2+^ concentrations were quantified by HPLC. Data are mean ± s.d. for *n* = 3 independent replicates and are representative of 3 independent experiments. **d**, Comparison of the activities of cGAS and its D84A/D86A mutant. Data are representative of 3 independent experiments. **e**, cGAS product was degraded by nuclease P1 but not CIP, as analyzed by HPLC. Data are representative of 3 independent experiments. **f**, Comparison of the retention time of possible cGAMP variants (3′,3′-cGAMP, 2′,3′-cGAMP, and 3′,2′-cGAMP) and the cGAS product. Data are representative of 3 independent experiments. **g**, MS/MS fragmentation spectra of the cGAS product (top) and the standard 3′,2′-cGAMP (bottom), and the chemical structure of 3′,2′-cGAMP. **h**, Michaelis-Menten kinetics plot of ATP and GTP for cGAS. Data are mean ± s.d. for *n* = 3 independent replicates and are representative of 3 independent experiments.

**Extended Data Fig. 2.**
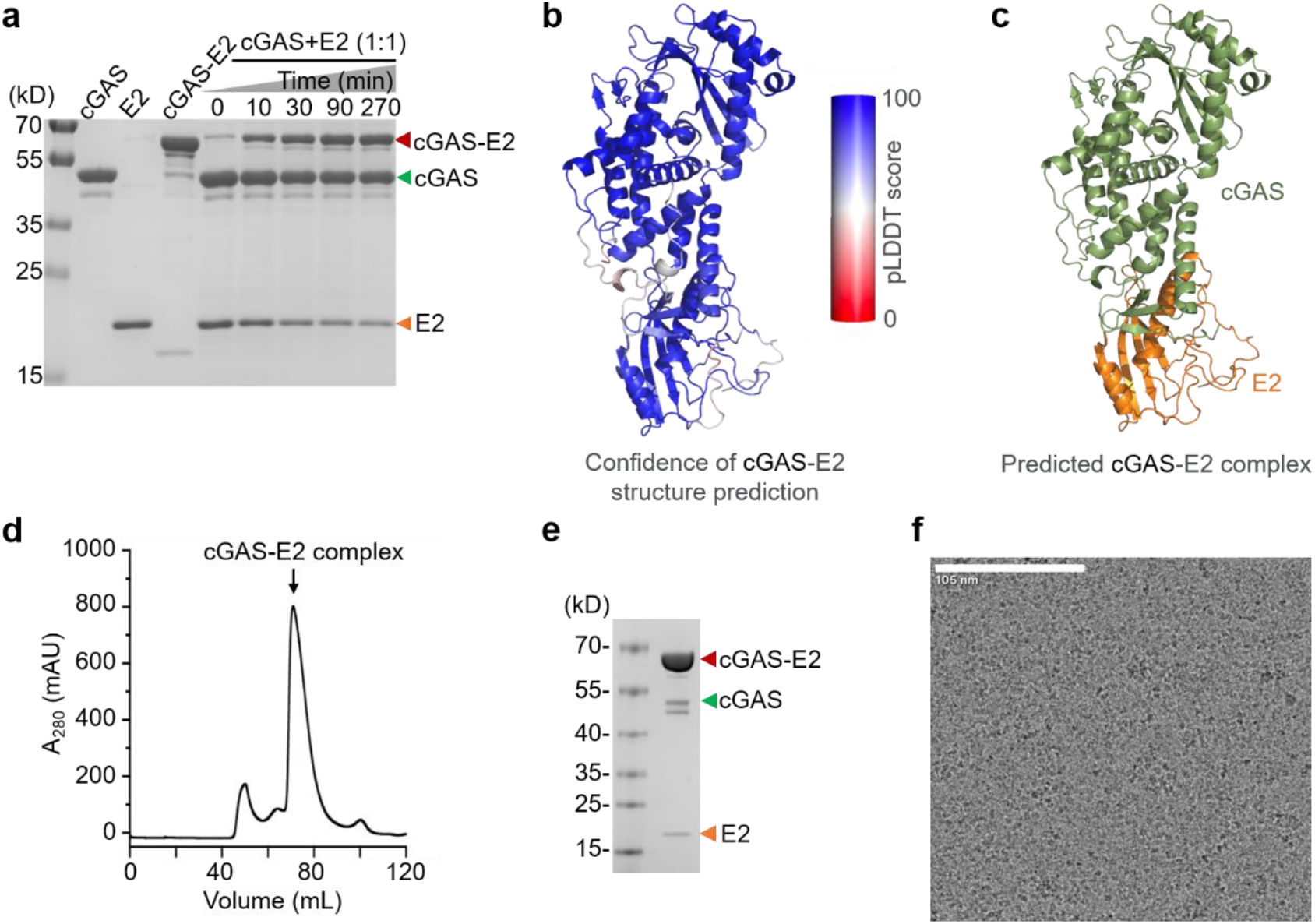
Predicted structure of the cGAS-E2 complex. **a**, SDS-PAGE analysis of the *in vitro* assembly of the covalent cGAS-E2 protein. Two proteins were mixed at a molar ratio of 1:1 and incubated at 4 °C for the indicated periods (cGAS+E2). Since both cGAS and E2 were His-tagged during the *in vitro* reconstruction, the molecular weight of cGAS-E2 protein was slightly higher than that obtained *in vivo* from His-tagged cGAS and non-tagged E2. Data are representative of 3 independent experiments. **b**, Ribbon diagram of the top ranked structure prediction of cGAS-E2 complex colored by the per-residue LDDT scores. **c**, Ribbon diagram of the top ranked structure prediction of cGAS-E2 complex with cGAS colored in green and E2 colored in orange. **d-e**, Chromatography traces and SDS-PAGE analysis of purified cGAS-E2 complex. **f**, Representative cryo-EM micrograph of cGAS-E2 complex.

**Extended Data Fig. 3.**
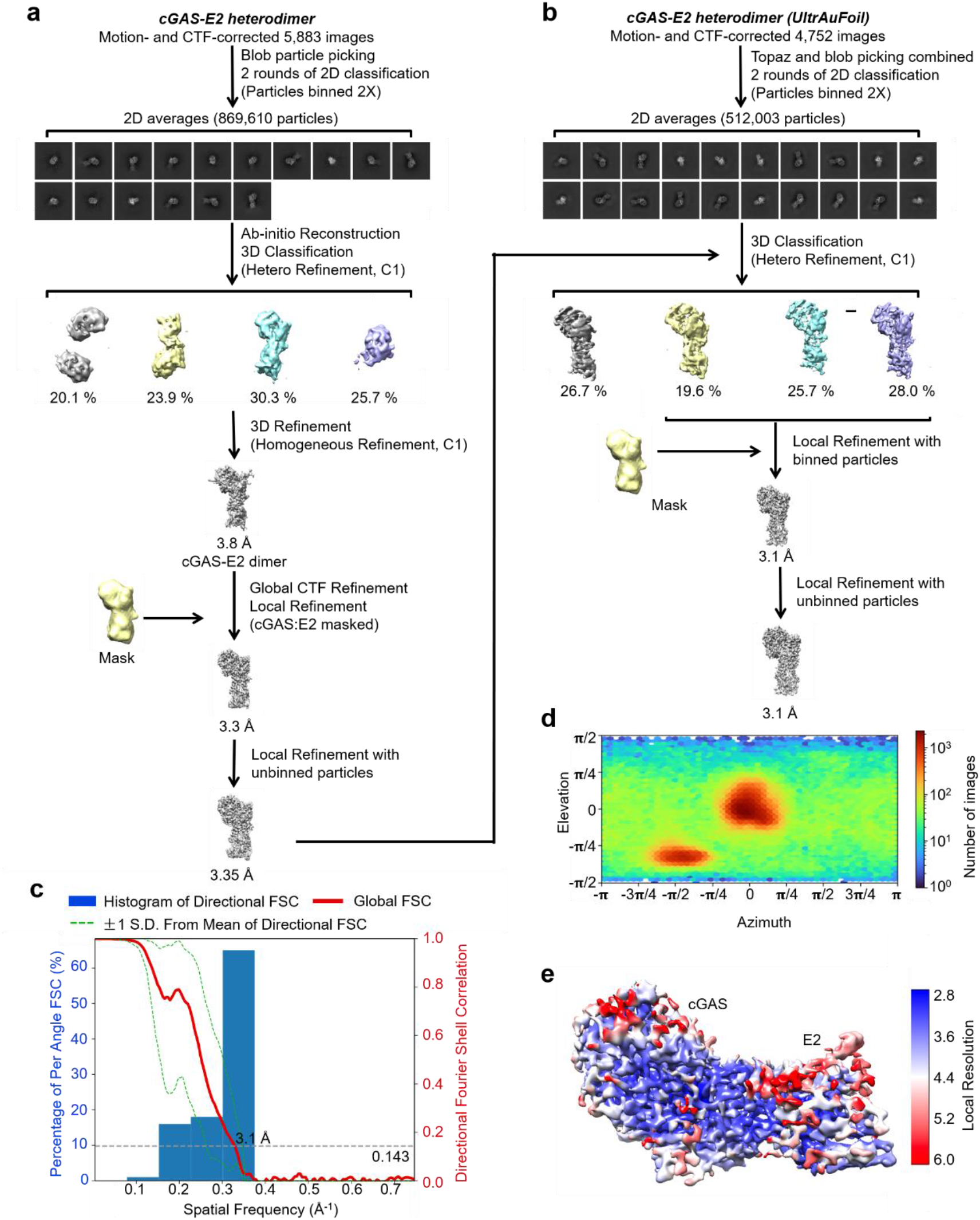
Cryo-EM workflows and the quality of reconstructed cryo-EM maps. **a-b**, Workflow of 3D reconstruction of the initial cGAS-E2 dataset (**a**) and the UltrAuFoil cGAS-E2 dataset (**b**). **c**, Directional FSC of the cGAS-E2 cryo-EM density map. **d**, Orientation distributions of cGAS-E2 3D reconstruction. **e,** Local resolutions of the cGAS-E2 cryo-EM map.

**Extended Data Fig. 4.**
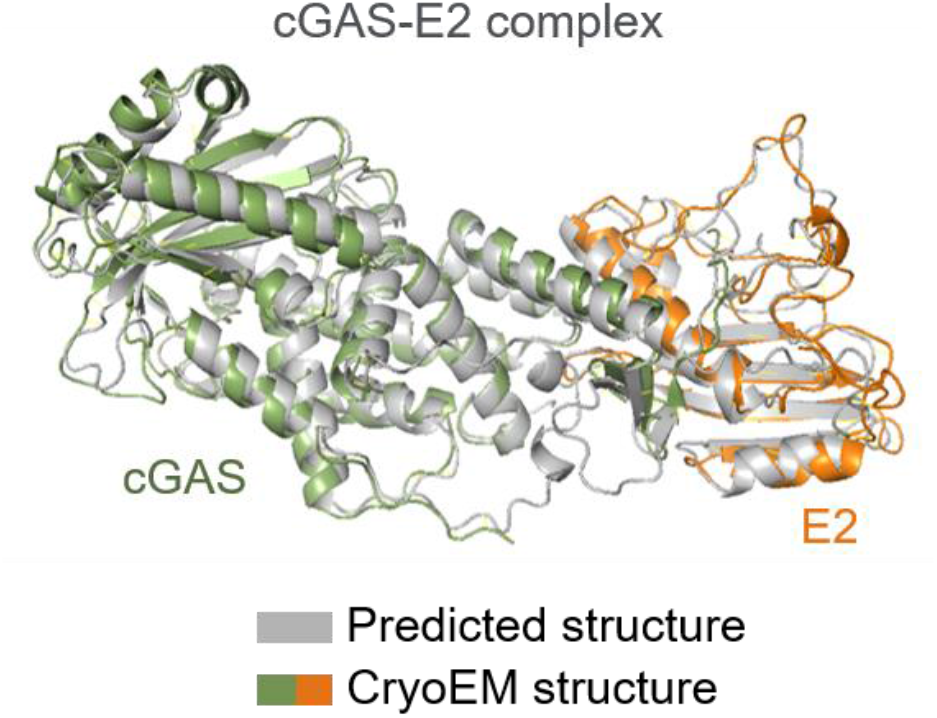
Structural comparison between experimental and predicted cGAS-E2. Superimposed predicted and experimental structures of cGAS-E2.

**Extended Data Fig. 5.**
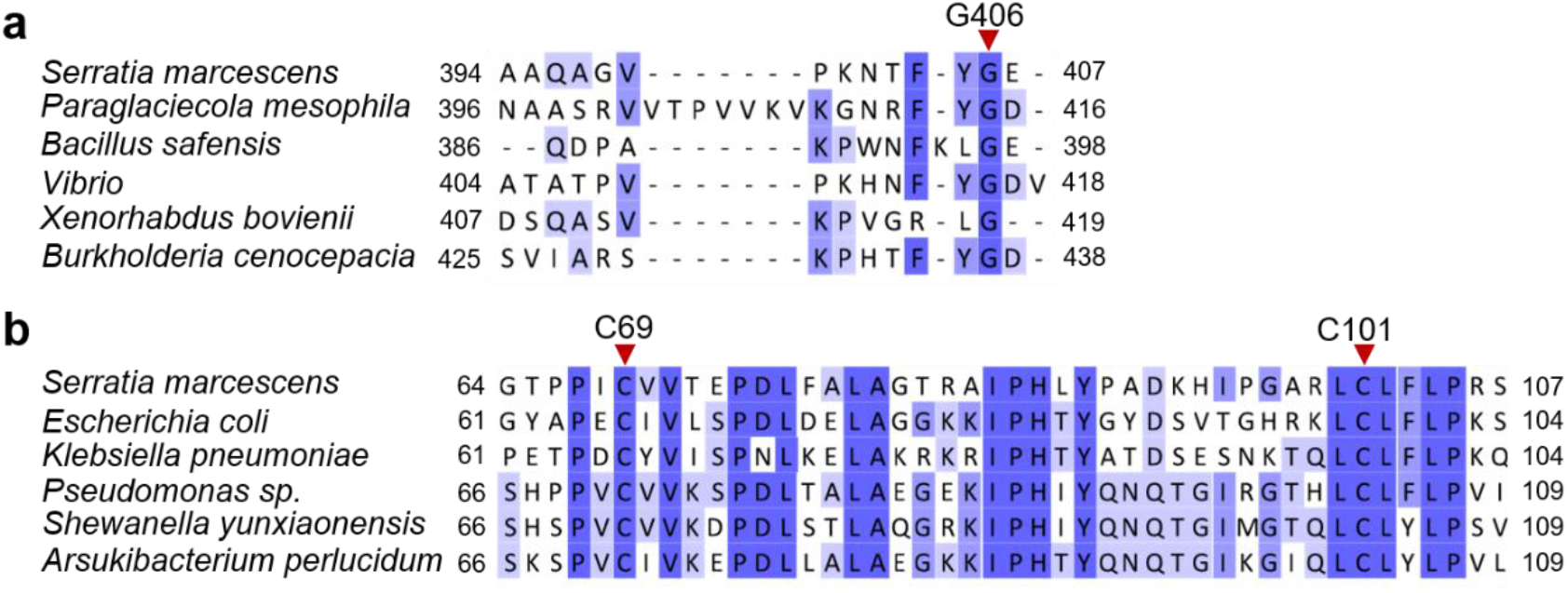
Sequence alignment of cGAS and E2 homologs. **a**, Sequence alignment of the C-terminal residues of cGAS and CD-NTase homologs from other E2-CBASSs. A conserved C-terminal glycine is marked by a red triangle. **b**, Sequence alignment of E2 and its orthologs shows two conserved cysteine residues (marked by red triangles).

**Extended Data Fig. 6.**
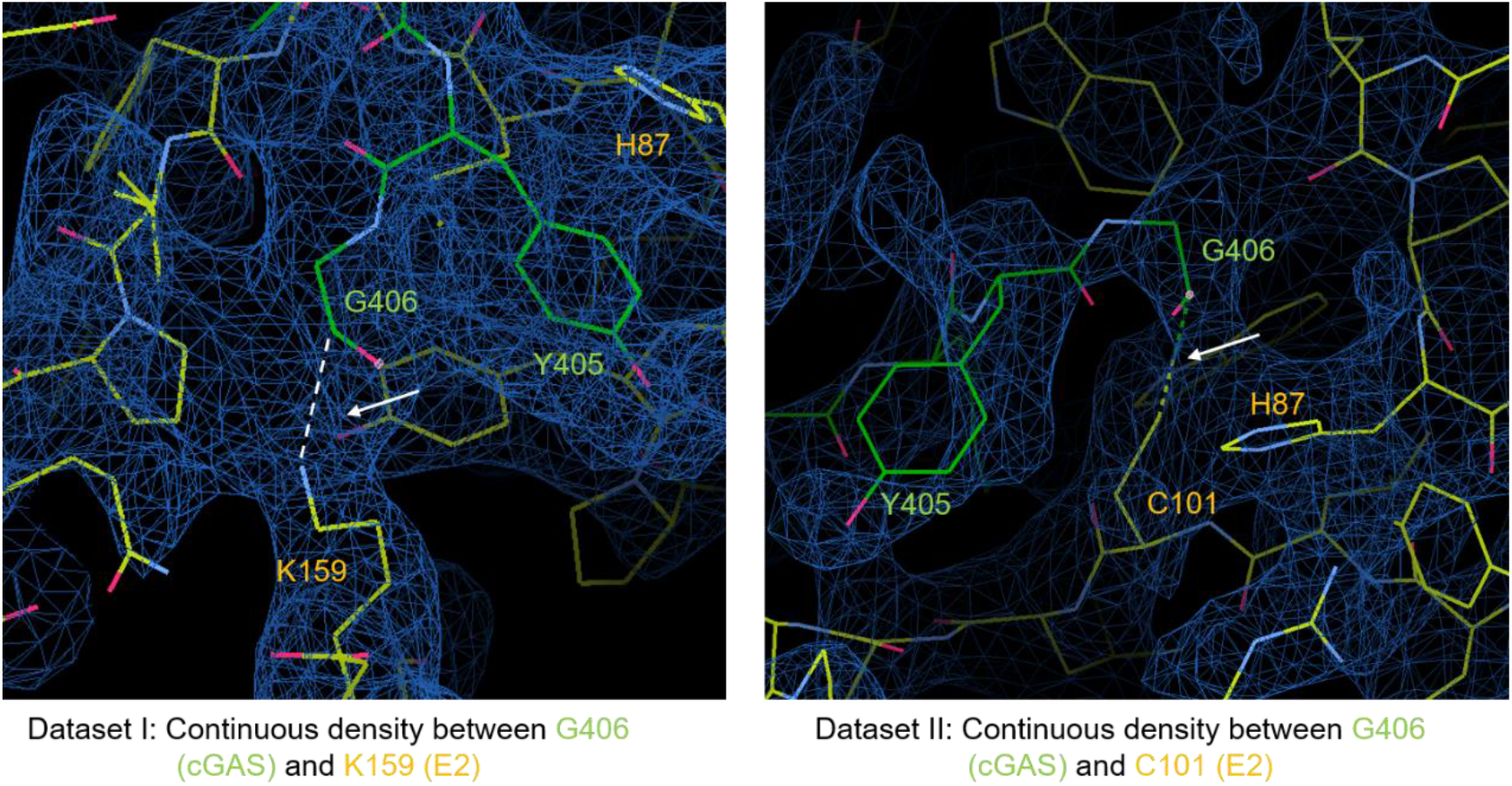
Magnified view of the E2 active site in cGAS-E2 complex, showing continuous density between G406 of cGAS and K159 or C101 of E2.

**Extended Data Fig. 7.**
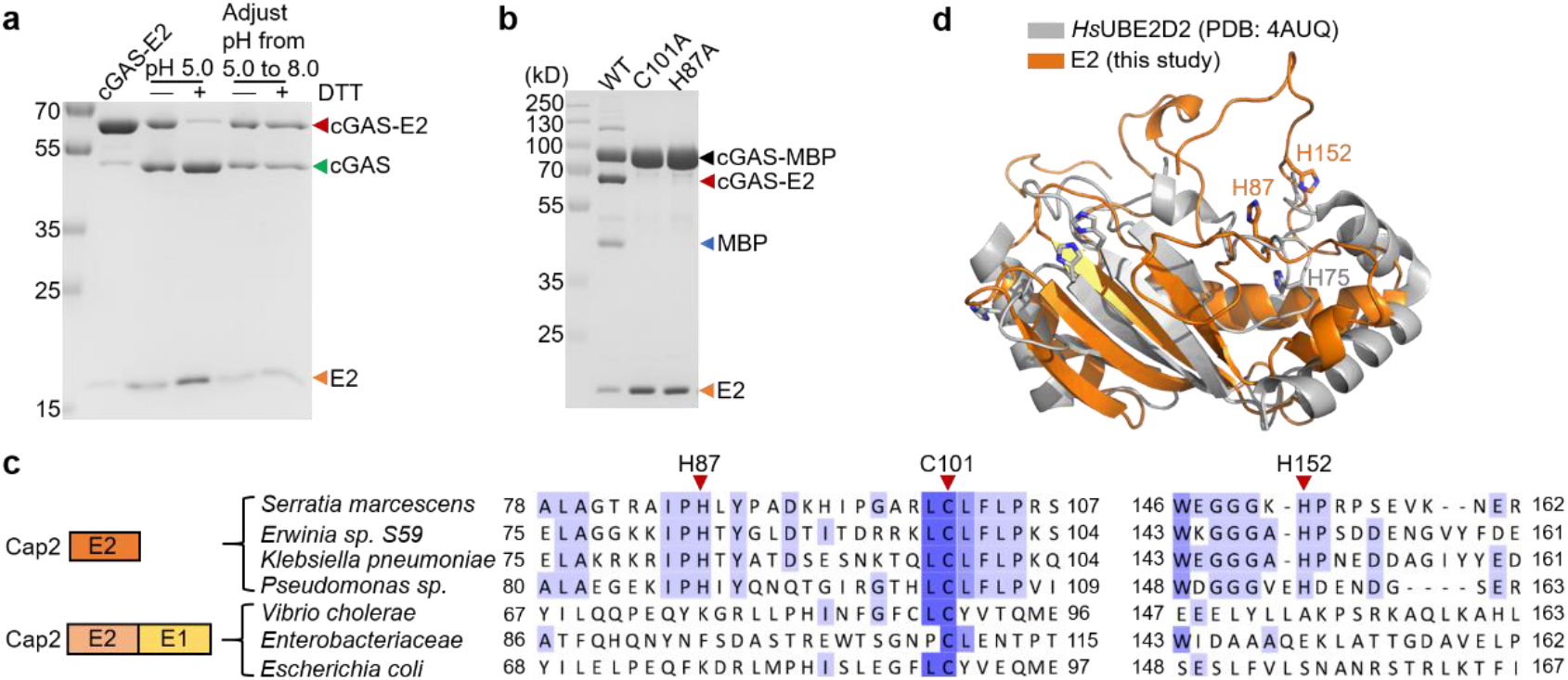
Two crucial histidine residues in E2 active site. **a**, SDS-PAGE analysis showed that the thioester bond (-DTT) and isopeptide bond (+DTT) in cGAS-E2 are interconvertible at various pH. Data are representative of 3 independent experiments. **b**, SDS-PAGE analysis of the *in vitro* processing of cGAS-MBP by E2 or E2 (C101A) or E2 (H87A). **c**, Sequence alignment of E2 homologs from E2-CBASSs and E1E2/JAB-CBASSs showed that they shared low homology. The two crucial histidine residues (marked by red triangles) are conservative in E2 from E2-CBASSs but not E1E2/JAB-CBASSs. **d**, Superimposed structures of E2 (orange) and human E2 (gray, PDB ID 4AUQ) showed that no similar catalytic site was found in human E2.

**Extended Data Fig. 8.**
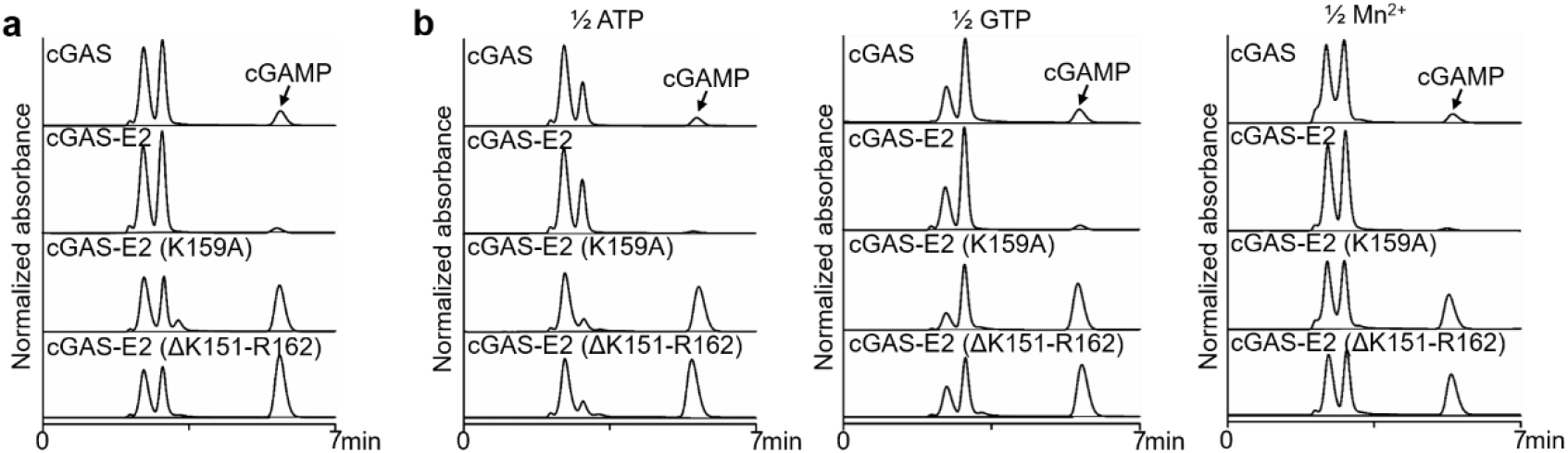
Regulation of cGAS activity by E2-mediated poly-cGASylation. **a**, HPLC analysis and quantification of the 3′,2′-cGAMP synthesized by cGAS and variants as shown in Fig. 4a. 20-μL standard reactions contained 800 nM enzyme, 250 μM ATP and 250 μM GTP in reaction buffer with 50 mM Tris-HCl (pH 7.5), 100 mM NaCl, 2.5 mM MnCl_2_, and 1 mM DTT. Reactions were incubated at 37 °C for 30 min, inactivated at 80 °C for 10 min, and centrifuged for 15 min at 21,000 × *g* to remove precipitated protein. **b**, HPLC analysis and quantification of the 3′,2′-cGAMP synthesized by cGAS and variants as shown in Fig. 4b, with concentration of ATP or GTP or Mn^2+^ in the reaction reduced by half compared to those in (**a**), respectively. Data are mean ± s.d. for *n* = 3 independent replicates and are representative of 3 independent experiments.

## Extended Data Methods

### Multiple Sequence Alignments

Alignment of cGAS or E2 homologs was performed using Clustal Omega and Jalview version 2.11.2.5. Matrix, BLOSUM; Open gap penalty, 10; Extending gap penalty, 0.05; End gap penalty, 10; Separation gap penalty 0.05.

### Steady-state kinetic measurement

For steady-state kinetic measurements of cGAS, the standard 20 μL-reaction mixture included 50 mM Tris-HCl (pH 7.5), 2.5 mM MnCl_2_, 100 mM NaCl, 1 mM DTT, 1 μM cGAS, 1 mM ATP (or GTP), and various concentrations (15.6, 31.3, 62.5, 125, 250, 500 and 1,000 μM) of GTP (or ATP). For each nucleotide concentration, samples were incubated at 37 °C for 15 min. Reactions were immediately heat-inactivated at 80 °C for 10 min and the samples were centrifuged for 15 min at 17,000 × *g* to remove precipitated protein. Of each sample, 10 μL was injected and reactions were analyzed by HPLC, as detailed above. Absorbance units were converted to μmol/L by comparing to a standard curve from 7.8 μM to 1 mM of chemically-synthesized 3′,2′-cGAMP (Biolog Life Sciences). Data were fitted by linear regression, and non-linear curve fitting Michalis-Menten kinetics and allosteric sigmoidal were calculated using GraphPad Prism version 8.0.2.

### Mass spectrometry

To characterize the product of cGAS, we performed LC-MS/MS on a Thermo Scientific™ UltiMate™ 3000 system coupled to a Thermo Scientific Orbitrap LC/MS (Q Exactive), using a Thermo Hypersil GOLD C18 column (100 mm × 2.1 mm, 3 μm) maintained at 25 °C with a flow rate of 0.25 mL/min. The mobile phase consisted of methanol (A) and 20 mM ammonium acetate (B). The HPLC gradient was as follows: 0–7 min, 2% A; 7–12 min, 2–30% A. The column was re-equilibrated for 7 min at 2% A. Detection was performed in positive ionization mode using an electrospray ionization (ESI) source under the following parameters: spray voltage, 3.2 kV; sheath and auxiliary gas flow rates, 40 and 15 arbitrary units, respectively; max spray current, 100.00 µA; S-Lens RF Level, 50%; capillary temperature, 300 °C; probe heater temperature, 350 °C. Profile MS1 spectra were acquired with the following settings: mass resolution, 70,000; AGC volume, 3 × 10^6^; maximum IT, 100 ms; scan range, 300– 1,000 m/z. Acquisition of data-dependent MS/MS spectra was performed using collision-induced dissociation (CID) with the following settings: mass resolution, 17,500; AGC volume, 1 × 10^5^; maximum IT, 50 ms; loop count, 10; isolation window, 4.0 m/z; normalized collision energy, 20, 40, and 60 eV. Data are reported for the z = 1 acquisition for each indicated cyclic nucleotide. The chemical structures were drawn using ChemDraw version 19.0.

### Proteomics analysis

Proteomic analysis is performed by a commercial company vendor (Genecreate). The protein samples were mixed with NuPAGE LDS sample buffer (Invitrogen) and separated using NuPAGE 4 to 12%, Bis-Tris gel. The corresponding gel bands were then cut and washed with 40% H_2_O, 60% acetonitrile and 50 mM NH_4_HCO_3_ (pH 8.0), and the protein bands to be identified were reduced and alkylated using TCEP and chloroacetamide respectively. The gel pieces were digested with trypsin (20 ng/mL) in 50 mM NH_4_HCO_3_ (pH 8.0) at 37 °C overnight. The digested peptides were extracted with 40% H_2_O, 60% acetonitrile and 1% formic acid, dried and reconstituted with 1% formic acid for LC-MS/MS analysis. The tryptic peptides (100 ng) were analyzed on a UltiMate 3000 RSLCnano coupled to an Q Exactive HF mass spectrometer (Thermo Fisher Scientific). The peptides were firstly loaded onto a 75 μm×2 cm trap column and separated on a 75 μm×25 cm Pepmap C18 analytical column (Thermo Fisher Scientific) with a binary buffer system. Buffer A was 0.1% formic acid and 3% DMSO in 100% H_2_O and buffer B was 0.1% formic acid and 3% DMSO in 80% acetonitrile with 20% H_2_O. The Eclipse mass spectrometer was operated in data-dependent acquisition mode with one full MS scan followed by MS/MS scans with higher-energy collision-induced dissociation (HCD) fragmentation. The typical MS settings were spray voltage of 2.2 kV, heated capillary temperature of 320 °C, HCD energy of 27. The LC-MS/MS RAW data were processed with Maxquant (version 2.1.2.0) for protein identification using a tailored database containing the sequences of E2 and cGAS and the *Escherichia coli* strain B/BL21-DE3 proteomes database in Uniprot. The typical search parameters were enzymatic specificity of Trypsin/P with maximum of two missed cleavages and variable modifications of oxidation on methionine. Carbamidomethyl (Cys) was set as the fixed modification. All results were screened at 1% protein and peptide level FDR. Only high confidence identified peptides were chosen for downstream protein identification analysis.

## Notes

### Competing Interest Statement

The authors have declared no competing interest.

### Summary of Updates

Mechanism and model revised based on additional results.

